# Quantitative 3D cytoarchitecture of human brain organoids using light-sheet microscopy

**DOI:** 10.64898/2026.03.19.712931

**Authors:** Omkar Suhas Vinchure, Amy Vivien Job, Hugo Alonso-Olivares, Fowzan S Alkuraya, Elke Gabriel, Jay Gopalakrishnan

## Abstract

Human brain organoids are valuable models for studying human brain development and disorders. Although organoids enable multi-omics analyses, understanding developmental mechanisms requires precise three-dimensional mapping of cell types and their dynamics within their cytoarchitecture. Traditional thin-section immunohistochemistry and imaging provide insights but disrupt spatial information. Light-sheet microscopy allows volumetric imaging of intact organoids, but methods and analytical algorithms remain technically challenging. Here, we introduce LUCID-org, an optimized and reproducible technique for **L**ight-sheet imaging of **U**nsectioned, **C**leared, **I**mmunolabeled, and **D**epth-resolved human brain **org**anoids. Coupled with machine-learning–based quantitative analysis, LUCID-org enables unbiased 3D characterization of cell-type composition and cytoarchitecture. We validated this approach using brain organoids derived from a patient with *CENPJ*-mutated microcephaly, revealing quantifiable defects in the ventricular zone (VZ), progenitor density, dynamics, cell-type variability, VZ lumen volume, and neuronal distribution compared with healthy controls. The LUCID-org pipeline is straightforward, taking approximately one week, significantly cheaper, non-toxic, and compatible with standard light-sheet microscopes. Therefore, the LUCID-org method could serve as a benchmark for comprehensive 3D analysis of human brain organoids.

## Introduction

Human brain organoids are powerful three-dimensional *in vitro* models derived from pluripotent stem cells that self-organize into structures that recapitulate features of early brain development ^1–3^. These include the organization of ventricular zone-like (VZ) and cortical progenitor populations, primitive neuronal layers, and the sensory structures associated with them, providing insight into the cellular dynamics underlying the early stages of human corticogenesis ^4–7^. Brain organoids help investigate neurodevelopmental disorders that are difficult to study in developing human brain tissue due to limited access to it ^8^. When differentiated from patient-derived iPSCs harboring mutations in genes critical for brain development, patient-specific brain organoids have revealed previously unexpected cellular mechanisms underlying brain malformations, such as microcephaly, lissencephaly, heterotopia, and a variety of neurogenetic disorders that affect the developing brain ^9^ ^10–12^.

Although brain organoids enable the generation of multi-omics data and subsequent analyses to identify molecular signatures^13^ of regulated development and disease progression, a full understanding of developmental mechanisms requires precise three-dimensional mapping of cell types and their spatial arrangement within the complex 3D cytoarchitecture. Conventional thin-section immunohistochemistry, while reliable and widely used, inherently lacks spatial context. Practical limitations include labor-intensive, time-consuming procedures and the potential for sampling bias arising from two-dimensional sectioning of a three-dimensional structure.

Recent advances in light-sheet (LS) fluorescence microscopy now enable rapid 3D imaging of intact tissues with less photobleaching, making it a promising method for analyzing intact organoids^14–17^. Improved tissue-clearing techniques, combined with light-sheet imaging, have begun to reveal detailed structural features in biological tissues and organoids, enabling subcellular-resolution 3D phenotyping^18–21^. However, broader application has been limited by technical issues related to sample preparation, tissue opacity, antibody penetration, variability in clearing and imaging techniques, and, most importantly, the lack of algorithms for quantification^22,23^. Additionally, there is a need to develop standardized, reproducible workflows that integrate immunostaining, clearing, volumetric imaging, and quantitative analysis within reasonable timeframes to address key questions in human brain organoid research.

We developed an optimized, step-by-step, reproducible workflow for whole-organoid immunostaining, tissue clearing, and volumetric imaging of intact brain organoids using light-sheet microscopy. By integrating machine-learning–based quantitative analysis, our platform enables unbiased characterization of cell-type composition and cytoarchitectural organization in 3D. Using brain organoids derived from a patient with *CENPJ*-mutated microcephaly, we validate our approach’s ability to detect quantifiable developmental abnormalities. The described method and its application may serve as a standard for comprehensive three-dimensional quantitative analysis of human brain organoids and disease modeling.

### Development of LUCID-org workflow

#### Immunostaining of unsectioned brain organoids

While using brain organoids as a model system, we identified the need to develop a reliable immunostaining protocol for whole organoids. This is especially important for examining spatial details, such as cytoarchitecture, cell shapes, and organization in the VZ, as well as large structures, such as the VZ lumen, which are often disrupted or misinterpreted in traditional 2D thin-sections ^24^. Conversely, previously established whole-mount staining protocols rely on the use of sophisticated equipment, conjugated antibodies, longer incubation times, specialized reagent development, and complex sample handling ^25,26^. Our LUCID-org workflow focuses on the use of standard laboratory reagents, including primary and secondary antibodies. Additionally, our sodium dodecyl sulfate (SDS)-based permeabilization buffer and fish-gelatin-based blocking buffer enhance antibody penetration throughout the organoid and facilitate antigen retrieval, shorten incubation times, and reduce background noise. It is also compatible with a wide range of antibodies and probes used in routine organoid staining.

#### Tissue clearing of brain organoids

Handling transparent tissues for imaging is challenging. We created a simple custom setup to hold the organoids in 3D within an agar column using an insulin syringe, which protects the immunostained organoids while allowing easy handling for repeated use and long-term storage. Initially, we used commercially available tissue glue to mount the sample, but it was detrimental to tissue clearing, rendering the sample permanently opaque and unsuitable for imaging. Our custom-made mounting assembly, which includes a cut syringe with a needle hook, greatly simplifies mounting and locating the organoids for imaging under a microscope, a major challenge when working with small, cleared samples (less than 2 mm). Moreover, embedding the organoids in an agar column enables rapid, non-toxic ethyl cinnamate-based tissue clearing. We present an efficient tissue-clearing protocol emphasizing simplicity, cost-effectiveness, ease of handling, and sample reuse for imaging.

#### Overview of the procedure

Our LUCID-org workflow is designed for quantitative developmental biology in brain organoids derived from iPSCs. It involves preparing samples through fixing in 4% paraformaldehyde, permeabilization, antigen retrieval, and blocking steps (Steps 1-10, Phase 1), whole-mount staining targeting specific antigens (Steps 11-18, Phase 2), embedding and clearing samples (Steps 19-38, Phase 3), followed by mounting for volumetric imaging (Steps 39-56, Phase 4). Additionally, the workflow is integrated to quantify data. As described below, we also highlight the development and use of our final sample assembly for mounting and reliable imaging on the light-sheet microscope.

#### Applications of the method

The LUCID-org workflow is suitable for imaging intact, stained, and tissue-cleared brain organoids of any type. It includes an essential data analysis pipeline with machine learning algorithms and can be easily adapted to different biological samples, such as organoids, assembloids, tumor spheroids, cell clusters, and larger samples like neonatal mouse brains, with minor adjustments to incubation times.

For human brain organoids, our approach can analyze and clarify organoid volumes, VZ organization, and the lumen, as well as microscopic features such as cell types and their spatial distribution, including cellular organelles like primary cilia, which are essential for studying the developmental mechanisms of brain tissues ^27^ ^28^. We validated our approach by modeling microcephaly using brain organoids derived from a patient with microcephaly and a *CENPJ mutation*, a centrosomal gene ^10,29^. Overall, the LUCID-org workflow can be integrated into a laboratory setting and used to address a wide range of biological questions, including high-throughput disease modeling.

#### Comparison with other methods

Protocols focused on tissue staining and clearing have been previously established which were primarily designed based on tissue type, size, multiplexing, and imaging modality, with particular attention to the extent of clearing and architectural preservation ^18–21^. The CLARITY protocol involves hydrogel-embedding of samples prior to staining ^19,30^, whereas the iDISCO protocol uses methanol dehydration and bleaching ^18,21^.

Our LUCID-org fixes tissues in a warm solution of 4% paraformaldehyde, a common laboratory fixative. Like other protocols, the LUCID-org approach also considers antibody multiplexing. While other protocols require longer incubation times for primary and secondary antibody diffusion (approximately 4 days), the passive diffusion-based staining in the LUCID-org workflow takes only a few hours to overnight, making it significantly faster. Additionally, it can be used for medium- to large-sized tissue samples, similar to the CUBIC ^20,25^ and CLARITY ^19,30^ protocols.

The LUCID-org offers high optical transparency similar to protocols like CLARITY, CUBIC, and iDISCO^18–21,30^. It uses a safer clearing agent, Ethyl Cinnamate (ECi), compared to iDISCO and CLARITY, which rely on more toxic agents such as dibenzyl ether (DBE) and acrylamide^19,21^. The clearing process in the LUCID-org workflow is notably faster, taking about 2 days versus around 7 days or more with other methods. We achieved high-quality staining while preserving tissue structure in human brain organoids. Overall, the LUCID-org workflow provides a cost-effective, quick, and safe solution for whole-mount staining and clearing, enhancing compatibility with light-sheet microscopy and other protocols. A detailed multi-parametric comparison of CLARITY, CUBIC, iDISCO, and LUCID-org is presented below (**Tables 1 and 2**).

**Table 1.** Comparison of immunostaining parameters.

| Parameter | CLARITY | CUBIC | iDISCO/iDISCO+ | LUCID-org |
| --- | --- | --- | --- | --- |
| Tissue preparation | Hydrogel-embedding | CUBIC-1 treatment | Methanol-dehydration and bleaching | PFA fixation |
| Permeabilization strategy | SDS lipid removal + Triton X-100 | Amino-alcohol delipidation + Triton X-100 | Strong solvent permeabilization (methanol + detergents) | SDS+Triton X-100 |
| Blocking agent | 3–10% normal serum (donkey/goat) + BSA + Triton X-100 | BSA or serum + Triton X-100 | Serum + DMSO + Triton X-100 | Fish gelatin+Triton X-100 |
| Primary antibody type | Standard unconjugated | Standard unconjugated | Standard unconjugated | Standard unconjugated |
| Secondary antibody type | Fluorophore-conjugated | Fluorophore-conjugated | Fluorophore-conjugated | Fluorophore-conjugated |
| Primary antibody incubation | 2–5 days | 1–4 days | 1–3 days | 4 hours to overnight |
| Secondary antibody incubation | 1–3 days | 1–2 days | 1–2 days | Overnight |
| Total staining duration | ~3–7 days | ~2–5 days | ~2–4 days | 2 days |
| Temperature during staining | 37 °C or RT | RT | RT | RT |
| Agitation requirement | Gentle shaking | Gentle shaking | Continuous shaking | Gentle rocking |
| Antibody penetration depth | High | Moderate | High | Comparable to iDISCO/iDISCO+ |
| Signal-to-noise ratio | Good | Good | Good | Comparable to iDISCO/iDISCO+ |
| Multiplex staining | Good | Moderate | Moderate | Comparable to CLARITY |
| Risk of epitope damage | Low | Low | Moderate | Low |
| Typical antibody concentration used | Standard (1:100–1:500) | Standard (1:100–1:500) | Often higher (1:50–1:200) | Standard |
| Overall staining efficacy | High | Moderate-High | High | High |

**Table 2.** Comparison of tissue clearing parameters.

| Parameter | CLARITY | CUBIC | iDISCO / iDISCO+ | LUCID-org |
| --- | --- | --- | --- | --- |
| Clearing principle | Hydrogel embedding + lipid removal | Hyperhydration and delipidation | Organic solvent dehydration and refractive index matching | Ethanol dehydration+refractive index matching |
| Clearing agent | Acrylamide hydrogel + SDS detergent | CUBIC-1 (urea, amino alcohols, Triton X-100) and CUBIC-2 RI solution | Methanol series, dichloromethane (DCM), dibenzyl ether (DBE) | Ethyl Cinnamate (ECi) |
| Clearing duration | ~5–14 days | ~3–10 days | ~1–3 days | 3 days |
| Optical transparency | High | Moderate | High | Comparable to CLARITY |
| Refractive index matching solution | Histodenz | CUBIC-2 or CUBIC-R+ | DBE | ECi |
| Tissue size | Medium–large tissues | Small–medium tissues | Medium–very large tissues | Small-medium tissues |
| Morphological changes in sample | Minimal distortion | Mild swelling (tissue expansion possible) | Tissue shrinkage common | Minimal distortion |
| 3D Structural preservation | Good | Good | Moderate | Comparable to CLARITY |
| Compatibility with light-sheet microscopy | Good | Good | Good | Comparable to CLARITY |
| Compatibility with confocal microscopy | Good | Good | Moderate | Comparable to CLARITY |
| Hazardous chemicals | Acrylamide (neurotoxic), SDS | Low toxicity aqueous chemicals | High toxicity organic solvents | None |
| Risk to microscope objectives | Low | Low | High if solvent contacts optics | Low |
| Cost effectiveness | Moderate | Moderate | High | Low |
| Ease of implementation | Moderate | Easy | Moderate | Easy |

#### Limitations of this study

Our study has certain limitations. Like any other method, our approach may be constrained by antibody compatibility, likely due to limited antibody penetration. These issues highlight the need to further optimize antibody selection, labeling techniques, and staining protocols for volumetric imaging. Our analysis was limited to a few developmental time points and focused mainly on a single genetic microcephaly model caused by a *CENPJ* mutation ^10,29^. Expanding imaging and analysis to additional developmental stages, genetic backgrounds, and disease models will be essential to evaluate the generalizability and robustness of this approach.

### Experimental design Brain organoid generation

#### hiPSCs thawing and maintenance

Human induced pluripotent stem cells (hiPSCs) were thawed from the cryobank and seeded onto Matrigel-coated 6 cm tissue culture dishes. Cells were maintained at 37 °C and 5% CO₂ in mTeSR™ Plus medium supplemented with 10 µM ROCK inhibitor (ROCKi) and 1% Penicillin/Streptomycin (PS) for the first 24–48 hours. Subsequently, half-medium changes were performed daily based on cell consumption, using mTeSR™ Plus with PS, while colony morphology was closely monitored. For expansion, cells were passaged at a 1:4 ratio using the enzyme-free dissociation reagent ReLeSR™ and reseeded onto fresh Matrigel-coated dishes following the manufacturer’s protocol.

#### Neural induction and neurosphere formation

We generated Hi-Q brain organoids using our recently published protocol^31^. Briefly, microwell plates were coated with anti-adherent Biofloat, centrifuged, incubated, rinsed with DMEM/F12, and filled with Neural Induction Medium (NIM) supplemented with ROCK inhibitor (ROCKi). hiPSC colonies were dissociated with Accutase, collected, centrifuged, and resuspended in NIM plus ROCKi. Cells were counted, viability assessed, and adjusted to 3 × 10⁶ cells/ml. One milliliter of cell suspension was seeded into each microwell containing 1 ml of NIM plus ROCKi, gently mixed, sealed, and centrifuged twice to ensure uniform distribution (∼10,000 cells per microwell). Plates were incubated overnight (Day 0) at 37 °C and 5% CO₂. Over the next 5 days, medium was partially replaced daily with fresh NIM without ROCKi to promote aggregation and neurosphere formation.

#### Organoid differentiation and maturation

Neurosphere transfer was performed using 250 ml spinner flasks pre-coated with Sigmacote to prevent adhesion. Flasks were filled with 80 ml of organoid differentiation medium supplemented with SB431542 and Dorsomorphin, then equilibrated on a rotating platform. Neurospheres were detached by gentle pipetting with a cut 1 ml tip, collected on a 40 µm strainer, and residual spheres recovered with DMEM/F12. Spheres were flushed into a suspension dish, collected, and transferred into spinner flasks. Cultures were maintained at 37 °C and 5% CO₂ with bi-directional rotation at 25 rpm. Medium was refreshed weekly by replacing 20 ml with fresh differentiation medium. After 30 days, cultures were switched to maturation medium lacking SMAD inhibitors and maintained until harvest at defined time points.

#### Phase 1: Human brain organoid fixation, permeabilization, and blocking (Figure 1) (Steps 1-10)

Organoids were washed three times with 1x phosphate-buffered saline (PBS) in 2 ml round-bottom microcentrifuge tubes. They were then fixed in 700 µl of pre-warmed 4% paraformaldehyde (PFA) for 1 hour at room temperature (RT). Afterward, samples were washed three times with 1x PBS to remove any residual PFA and stored in PBS at 4 °C. For immunostaining, PBS was removed, and organoids were incubated in 700 µl permeabilization buffer for 1 hour at RT on a rocking platform, followed by three washes with 1x PBS. Subsequently, organoids were incubated in 700 µl blocking buffer for 1.5 hours at RT with gentle rocking (**Figure 1A**).

**Figure 1:**
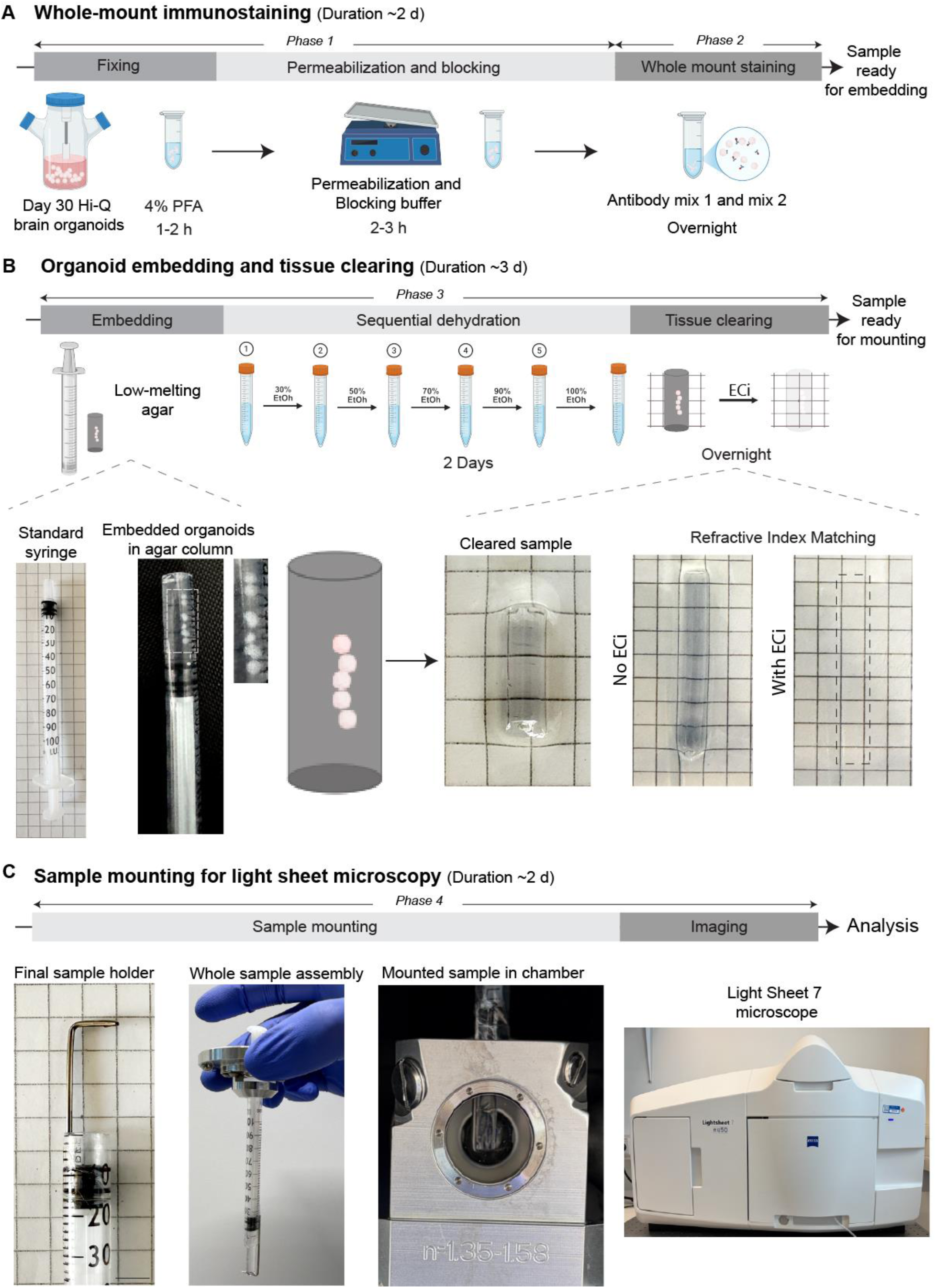
Steps for sample preparation for light sheet fluorescence microscopy, including whole-mount staining of brain organoids, embedding, tissue clearing, and mounting. **A**. Overview for the whole-mount staining protocol for immune-labelling of brain organoids for LS fluorescence microscopy with organoid fixation, permeabilization, and blocking, and multiplexed staining with a cocktail of primary and secondary antibodies. **B.** Overview of embedding organoids in an agar column using a custom-made syringe assembly, followed by dehydration steps for tissue clearing and refractive index matching with Ethyl Cinnamate as the clearing agent. The inset in the left panel shows stacked organoids within the agar column. Insets on the right show that the refractive index matches that of the agar column containing brain organoids, which is now completely transparent. Dotted lines indicate the position of the agar column. The scale bar on the grid represents 5 mm. **C.** Custom-made holder and final sample assembly for mounting in the sample chamber on the LS7 microscope.

#### Phase 2: Whole-mount immunostaining of brain organoids (Figure 1) (Steps 11-18)

For primary antibody staining, Antibody Mix 1 (Mix 1) and Antibody Mix 2 were prepared separately in blocking buffer (**Table 3**) and supplemented with 2% Sodium azide for repeated use. Organoids were incubated in 700 µl of the respective primary antibody mixture overnight at room temperature on a rocking platform. The following day, primary antibody solutions were retrieved, and organoids were washed three times for 20 minutes each in blocking buffer. For ciliary staining, organoids were then incubated with the ARL13B primary antibody in blocking buffer for 3–4 hours at room temperature on a rocking platform, followed by three 20-minute washes with 1x PBS. Secondary antibodies conjugated to Alexa Fluor dyes (**Table 4**) were prepared in blocking buffer, and organoids were incubated in 700 µl of the secondary antibody solution overnight at room temperature with gentle rocking, protected from light. The next day, samples were washed three times for 20 minutes each in blocking buffer, with the final wash extended up to 3 hours to reduce fluorescence background (**Figure 1A**). Organoids were then ready for agarose embedding.

**Table 3.** Antibody panel used in this study.

| Cell line | Condition | Antibodies |
| --- | --- | --- |
| Control | Mix 1 | α-SOX2, α-MAP2, α-ARL13B |
| Control | Mix 2 | α-PTPRZ1, α-p-Vimentin, α-ARL13B |
| *CENPJ* | Mix 1 | α-SOX2, α-MAP2, α-ARL13B |
| *CENPJ* | Mix 2 | α-PTPRZ1, α-p-Vimentin, α-ARL13B |

**Table 4.** Antibodies.

| Type | Name | Host | Dilution | Company | Catalog no. |
| --- | --- | --- | --- | --- | --- |
| Primary | α-SOX2 | Mouse | 1:400 | Abcam | ab79351 |
| Primary | α-MAP2 | Rabbit | 1:400 | ProteinTech | 17490-1-AP |
| Primary | α-ARL13B | Rat | 1:400 | BiCell Scientific | 90413h |
| Primary | α-PTPRZ1 | Rabbit | 1:400 | Sigma | HPA015103 |
| Primary | α-p-Vimentin | Mouse | 1:400 | MBL | DO76-3 |
| Secondary | α-mouse 488 | Goat | 1:1500 | Sigma | 62197-1ML-F |
| Secondary | α-rabbit 594 | Goat | 1:1500 | Invitrogen | A23740 |
| Secondary | α-rat 647 | Chicken | 1:1500 | Invitrogen | A21472 |

**Table 5.** Buffer compositions.

| Name | Use for | Composition | Required concentration |
| --- | --- | --- | --- |
| Permeabilization buffer | Whole-mount Immunofluorescence staining | Glycine | 30 mM |
|  |  | Triton X-100 | 0.25% |
|  |  | Tween20 | 0.2% |
|  |  | SDS | 0.1% |
| Blocking buffer | Whole-mount Immunofluorescence staining | Glycine | 30 mM |
|  |  | Cold water fish gelatine | 0.05% |
|  |  | Triton X-100 | 0.2% |

#### Phase 3: Embedding, sequential dehydration, and tissue clearing of brain organoids (Figure 1) (Steps 19-38)

Organoids were embedded in 1.5% low-melting agarose prepared with deionized water. A 1 ml cut insulin syringe (without the tapered tip) was used for embedding. An agar bath was created by adding about 1.5 ml of molten agarose into a 2-ml round-bottom microcentrifuge tube, and the syringe was prefilled with approximately 0.4 ml of agarose. Organoids were transferred into the agar bath with minimal residual buffer and quickly aspirated into the syringe along with molten agarose, avoiding solidification. The organoids were spaced within the agar column, and the filled syringe was placed on a cooling pack while gently rotating until the agarose solidified. Then, the syringe was kept at 4°C for 15 minutes to ensure full solidification of the column. Subsequently, the agarose column was extruded into a 15-ml Falcon tube containing 12–14 ml of 30% ethanol and dehydrated on a rocking platform at room temperature through a graded series of ethanol solutions (30%, 50%, 70%, 90%, and absolute ethanol). Incubations lasted about 2 hours per step (or 4 hours in 90% ethanol), or alternatively, overnight at 4°C. After dehydration, ethanol was removed, and the samples were immersed in 12–14 ml of Ethyl Cinnamate (ECi) overnight without disturbance, ensuring no air bubbles formed. Once cleared, the column became transparent and sank in the tube (**Figure 1B**). The samples were then ready for mounting and imaging.

#### Phase 4: Sample mounting and imaging (Figure 2) (Steps 39-56)

For imaging, we used the Light Sheet 7 system. Objectives must match the refractive index of the clearing reagent Ethyl Cinnamate (RI 1.558). The appropriate sample chamber was mounted and filled with fresh ECi using a syringe and tubing (ECi can be reused up to 3–4 times). A custom assembly involving the syringe and a needle hook was prepared (**Figure 1C**). This hook supports the sample in space and holds it securely during imaging with Ethyl Cinnamate. A glass Petri dish was cleaned with distilled water, 70% ethanol, and absolute ethanol. The contents from the Falcon tube were transferred to the dish, and the column was gently lifted with metal forceps and inserted into the syringe. The piston was carefully pushed forward to position the column on the needle hook, and the syringe was mounted in the sample holder and placed in the microscope. Using the system software, the sample was adjusted so that the column faced the objective. Organoids were first located using brightfield and fluorescence overview imaging at 0.36x magnification in a 3-track configuration for independent laser adjustment.

**Figure 2:**
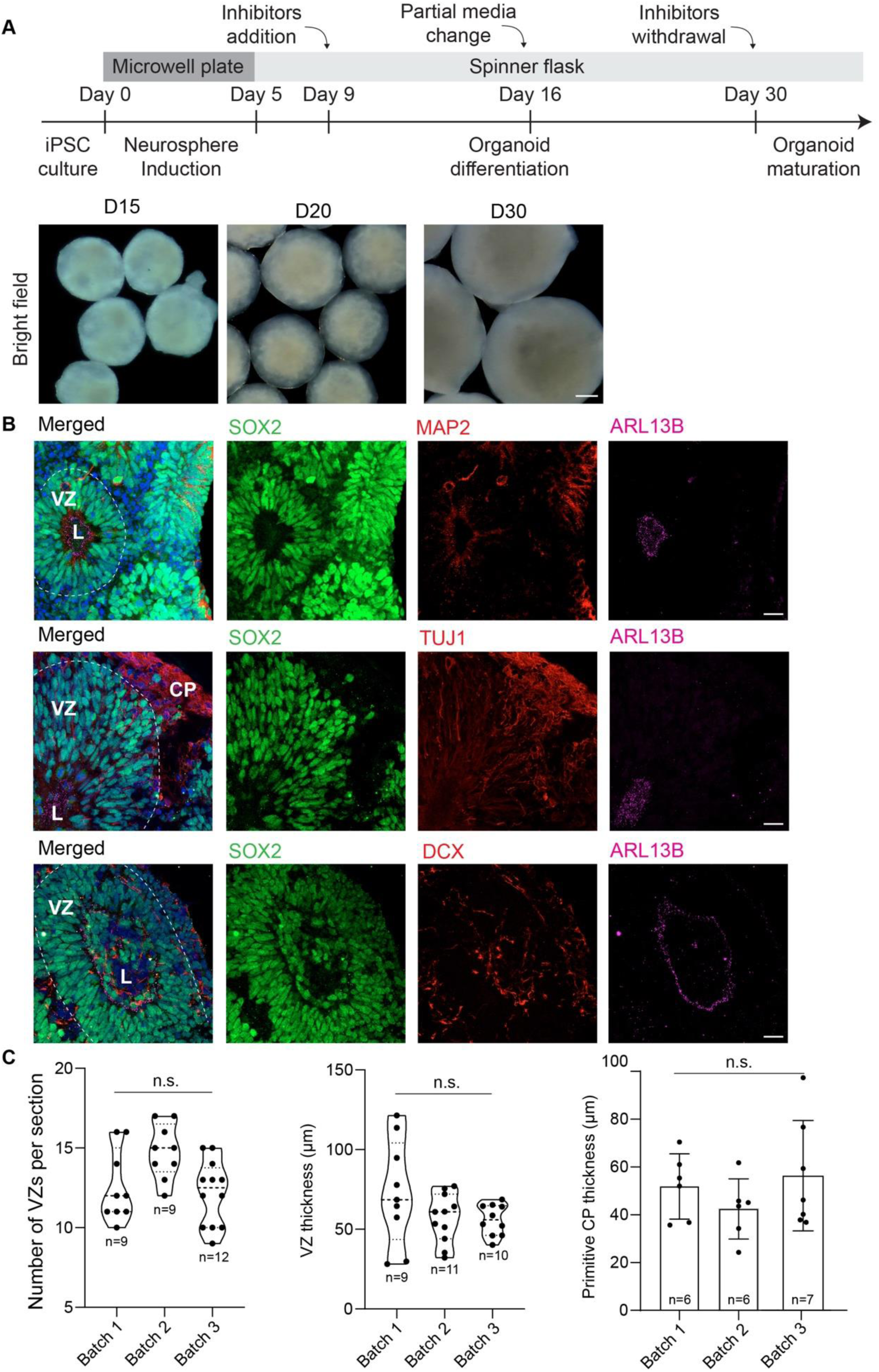
Conventional thin-sectioning and immunostaining of brain organoids reveal typical cytoarchitectures of developing brain tissue. **A**, Protocol overview for the generation of Hi-Q brain organoids from hiPSCs. Macroscopic images (bottom panel) at various time points. Scale bar, 200 μm. **B.** Thin-section and immunofluorescence staining of Day 30 brain organoids. ARL13B (magenta) specifies primary cilia extended into the ventricular lumen (L). SOX2 (green) labels apically organized NPCs forming a ventricular zone (VZ). Neuronal cells specified by MAP2, TUJ1, and DCX (red) are positioned at the basal side, forming a primitive cortical plate (CP). Scale bar, 20 μm. **C.** Quantifications of VZ numbers per section, VZ thickness, and primitive CP thickness. Error bars indicate standard deviation. The data are derived from at least 3 organoids per batch. The values for “n” and “p” are indicated in the plot. Statistical analysis was performed using one-way ANOVA. n.s. - not significant.

Imaging parameters such as laser power, exposure, magnification, detector size, and field of view were adjusted as needed. Pivot Scan was enabled to reduce shadow artifacts. Z-stack acquisitions were performed to image the entire organoid, and the data were saved to the processing computer. After imaging, the syringe was returned to the load position, and the sample holder was removed. The agarose column was retracted into the syringe and transferred into a labeled 15 ml Falcon tube containing ECi for storage. The system was then shut down, and Eci was recovered from the sample chamber for reuse, with all components rinsed using absolute ethanol. The sample chamber was air-dried prior to storage.

#### Expertise needed to implement the protocol

This procedure requires experience in cell culture, immunostaining, and imaging. Since the LUCID-ORG workflow emphasizes quantitative organoid biology, prior experience with organoids is recommended. Familiarity with fluorescence microscopy and 3D image analysis, using tools such as Arivis Pro, is also recommended.

## Materials

All the materials are readily available from certified laboratory material vendors.

### Biological material

A commercially available human iPSC cell line, IMR90, was used to generate control brain organoids. To model microcephaly, a patient-derived iPSC cell line with a *CENPJ* mutation ^10^ was used to generate *CENPJ*-associated microcephaly brain organoids.

### Reagents

#### Cell culture components

- DMEM/F12 (BenchStable™, ThermoFisher Scientific, Catalog # A4192002)
- mTeSR™ Plus medium (StemCell Technologies, Catalog # 100-0276)
- Neural induction medium (STEMdiff™, StemCell Technologies, Catalog # 05835)
- Penicillin-streptomycin (Gibco, Catalog # 15140-122)

▴ CAUTION: This mixture may cause allergic skin reactions and is suspected of harming fertility and the unborn child. Wear protective gloves, clothing, eye protection, and face protection.

- Neurobasal medium (Gibco, Catalog # 12348017)
- ReLeSR™ (StemCell Technologies, Catalog # 100-0483)
- Matrigel® Matrix (StemCell Technologies, Catalog # 354277)
- Accutase® (StemPro™, Gibco, Catalog # A11105-01)
- ROCK inhibitor (Y-27632 2HCl, Selleckchem, Catalog # S1049)

▴ CAUTION: Harmful if swallowed, in contact with skin, or inhaled. Wear protective gloves, clothing, eye protection, and face shield.

- Dorsomorphin (Selleckchem, Catalog # S7306)

▴ CAUTION: Harmful if swallowed. Very toxic to aquatic life with long-lasting effects. Avoid exposure to the environment.

- SB431542 (Selleckchem, Catalog # S1067)

▴ CAUTION: Harmful if swallowed. Very toxic to aquatic life with long-lasting effects. Avoid exposure to the environment.

- L-glutamine (ThermoFisher Scientific, Catalog # 25030081)
- MEM NEAA (Non-essential amino acids) (Gibco, Catalog # 11140-050)

▴ CAUTION: This substance can cause skin irritation and serious eye irritation. Wear protective gloves/protective clothing/eye protection/face protection.

- 2-mercaptoethanol (Gibco, Catalog # 31350-010)

▲ CAUTION: May cause an allergic skin reaction. Suspected of damaging fertility or the unborn child. Wear protective gloves/protective clothing/eye protection/face protection.

- N2 supplement (Gibco, Catalog # 17502-048)
- B27 supplement (Gibco, Catalog # 12587-001)
- Insulin solution human (Sigma-Aldrich, Catalog # I9278-5ML)
- Sigmacote® (Sigma-Aldrich, Catalog # SL2-100ML)

▲ CAUTION: This mixture is flammable and can cause skin and eye irritation. Work under the hood. Wear protective gloves/protective clothing/eye protection/face protection.

#### Fixation, permeabilization, and blocking

- Glycine (AppliChem, Catalog # A1067, 1000)
- Cold water fish gelatine (Sigma LifeScience, Catalog # G7041-500G)
- Triton™ X-100 (Sigma-Aldrich, Catalog # X100-500ML)

▲ CAUTION: Harmful if swallowed. Causes skin and serious eye irritation. Extremely toxic to aquatic life with long-lasting effects. May disrupt endocrine systems in the environment. Avoid releasing into the environment. Wear protective gloves, eye protection, and face protection.

- Tween®20 (Fisher Bioreagents, Catalog # BP337-500)
- PFA (Paraformaldehyde, Roth, Catalog # 0335.3)

▲ CAUTION: Flammable material. Can cause hazards such as skin, eye, and respiratory irritation. Work in a fume hood. Wear protective gloves and eye gear.

- PBS (Phosphate-buffered saline, Sigma-Aldrich, Catalog # P4417-100TAB)
- Sodium azide (ThermoScientific, Catalog # 190380050)

▲ CAUTION: Fatal if swallowed, in contact with skin, or inhaled. May cause organ damage (brain) through prolonged or repeated exposure if swallowed. Very toxic to aquatic life with long-lasting effects. Wear protective gloves and clothing. Avoid release into the environment.

- SDS (molecular biology grade) (Dodecyl Sulfate Sodium Salt, AppliChem, Catalog # A2263-0500)

Antibodies

For a list of antibodies, see Table 4

#### Embedding, tissue clearing, and imaging

- Ethyl Cinnamate (ECi, Sigma-Aldrich, Catalog # 8002380250)
- Low-melting agarose (Sigma-Aldrich, Catalog # A0701-25G)
- Absolute ethanol (molecular grade) (Roth, Catalog # 9065.1)

▴ CAUTION: Highly flammable liquid and vapour. Causes serious eye irritation. Work under a fume hood.

#### Equipment

- 15 and 50 mL Falcon tubes (Greiner BIO-ONE, Catalog # 188261-N, # 227270)
- 6 and 10 cm tissue culture dishes (SARSTEDT, Catalog # 83.3901, # 83.3902)
- 2, 5, 10, 25, and 50 ml serological pipettes (Greiner CELLSTAR®, Catalog # P7990, # P7615, # P7740, # P7865, # Z652555)
- 1,000, 200, 20, and 10 µl sterile filter pipette tips
- Serological pipette controller (accu-jet®S, BRAND, Catalog # 26351)
- Cell strainer (SARSTEDT, Catalog # 83.3945.040)
- 0.5, 1.5, and 2 ml Microcentrifuge tube (Eppendorf, Catalog # 0030121023, # 0030120086, # 0030120094)
- Hemocytometer, Neubauer improved (Superior Marienfeld, Catalog # 0640010)
- Scissors
- Cryotubes (Greiner BIO-ONE, Catalog # 126279)
- Microwell culture plate (AggreWell™800, Stemcell Technologies, Catalog # 34815)
- EVOS XL Core microscope (Invitrogen)
- Light sheet 7 microscope (ZEISS, custom)
- 250 ml Spinner flask (Pfeiffer, Catalog # 183 001)
- Magnetic platform (Pfeiffer, Catalog # 183 001)
- Dry bath (Thermo Scientific, Catalog # TSGP05)
- Cell culture cabinet/laminar flow (Thermo Scientific, HERASAFE 2030i)
- Rocker (Thermo Scientific, Catalog # 88882002)
- Centrifuge (Eppendorf, Centrifuge 5810)
- Incubator (Thermo Scientific, HERACELL VIOS 160i, HERACELL 150i)
- 1 ml insulin syringes (BRAUN, Catalog # 9161708V)
- Extension line (Type: Heidelberger, BRAUN, Catalog # 4097173)
- Razor blade
- Metal forceps
- Glass petri dish
- Needle
- Cooling pack
- Aluminium foil
- Parafilm
- Software: Arivis Pro, Zen Black, Zen Blue, MS package
- USB stick

#### Data analysis tools

- Zeiss Zen Black and Zen Blue software
- Zeiss Arivis Pro software
- GraphPad Prism

## Procedure

### Phase 1: Brain organoid fixation (Figure 1A)

1. Carefully transfer the organoids to a 2 ml round-bottom microcentrifuge tube with minimal pipetting to prevent structural damage. *Note:* Use a cut pipette tip to avoid mechanical harm to the organoids.
2. Rinse the organoids 3 times in PBS under a fume hood to remove residual medium.
3. Remove as much PBS as possible, then add 700 µl of warmed 4% Paraformaldehyde (PFA) to organoids. ▴ CAUTION: Work in a fume hood, as PFA is flammable and can cause skin, eye, and respiratory irritation. Wear protective gloves and eye protection.
4. Mount the tubes on a rocker platform at room temperature for 1 hour at 2-4 rpm (gentle mixing). ▴ CRITICAL STEP: Ensure the organoids can move freely in PFA inside the tube. Make sure they don’t clump together. Lightly tap the tube to help release the organoids.
5. Dispose of the PFA in the appropriate waste container under the fume hood, following proper waste disposal procedures.
6. Rinse the organoids three times in PBS. ▪ PAUSE POINT The fixed organoids can be stored at 4 °C in 2 ml of PBS for subsequent immunostaining. **Organoid permeabilization and blocking (Figure 1A)**
7. Carefully remove the PBS from the tube and add 700 µl of Permeabilization Buffer to the organoids (see reagents section for buffer compositions). ▲ CRITICAL STEP Ensure the Sodium Dodecyl Sulfate (SDS) detergent in the permeabilization buffer is completely dissolved by briefly warming the buffer to 37 °C.
8. Mount the tubes on the rocker platform for 1 hour at room temperature (gentle mixing). ▲ CRITICAL STEP For organoids with a diameter less than 1mm, perform the permeabilization step for 20 minutes. Ensure that the organoids can move freely within the PFA tube. Make sure they do not clump together. Lightly tap the tube to release the organoids and prevent bubble formation.
9. Remove the permeabilization buffer and wash the organoids three times for 5 minutes each in blocking buffer (see reagents section for buffer compositions; see Table 4).
10. Add 700 µl of fresh blocking buffer to the tubes. Place the tubes on the rocking platform at room temperature for 1.5 hours. **Phase 2: Whole-mount immunostaining (Figure 1A)**
11. Prepare the primary antibody mix in blocking buffer in a tube as follows. In this work, we used the following antibodies for Mix 1: Anti-SOX2 1:500; Anti-MAP2 1:500; anti-ARL13B 1:600, and for Mix 2: anti-PTPZR1 1:400, anti-p-Vimentin (Phospho-Vimentin) 1:400, and anti-ARL13B 1:600 (see Table 1, 2). ▲ CRITICAL: 2% sodium azide must be added to the primary antibody solutions to prevent bacterial contamination. This aids in long-term storage and repeated use.
12. Incubate the fixed and blocked organoids separately in the antibody solution overnight at room temperature on a rocking platform.
13. The next day, carefully retrieve the primary antibody mixture and wash the organoids three times for 20 minutes each with blocking buffer. ▲ CRITICAL STEP The anti-ARL13 B antibody is added separately in the protocol. Longer incubation times resulted in excessive background staining.
14. Incubate the organoids in anti-ARL13B solution for 3 to 4 hours at room temperature on a rocking platform.
15. Retrieve the antibody solution and wash the organoids three times for 20 minutes each using the blocking buffer.
16. Prepare the secondary antibody mix (see Table 1) in blocking buffer in the same tube as follows: goat anti-mouse 488 at 1:1500; donkey anti-rabbit 594 at 1:1500; chicken anti-rat 647 at 1:1500. ▲ CRITICAL: 4’,6-diamidino-2-phenylindole (DAPI) is incompatible with light sheet microscopy when using the clearing agent ethyl cinnamate. Alternative nuclear stains should be used if needed.
17. Cover the tubes with aluminum foil and incubate the organoids in the secondary antibody solution overnight at room temperature on the rocking platform.
18. The next day, wash the organoids 3 times for 20 minutes each with the blocking buffer. Extend the final wash step to 3 h at room temperature. ▴ CRITICAL STEP: This process removes the *‘*fluorescence halo*’* around the organoids, which interferes with optimal imaging. The organoids are now prepared for embedding in agarose. ▪ PAUSE POINT The stained organoids can be stored in PBS at 4 °C overnight, but it is recommended to proceed immediately with sample embedding to prevent bleaching and signal loss. **Phase 3: Brain organoid embedding and tissue clearing (Figure 1B) Embedding in agar:**
19. Prepare 1.5% low-melting agar in distilled water and warm it up till the solution is fully clear.
20. Allow the solution to cool until it is safe to touch. It is best to place the bottle on a heated platform while embedding.
21. In the meantime, prepare 1 mL insulin syringes by trimming the tapering tip with a razor blade until the barrel is cylindrical (**Figure 1B, first panel).**
22. Keep a cooling pack ready.
23. Make an agar bath by pouring 1.5ml molten agar into a 2ml round-bottom tube. Ideally, this agar bath could be kept warm in a heating block.
24. Depress the syringe piston to the 0.4 mL mark and pour molten agar into the syringe.
25. Using a cut pipette tip, transfer the organoids in a minimal volume of blocking buffer/PBS into the agar bath. ▲ CRITICAL STEP This step removes the wash solution surrounding the organoids and removes the surrounding *‘*envelope*’* that causes imaging issues.
26. With the agar, quickly transfer the organoids into the syringe and fill it to the brim. ▲ CRITICAL STEP: Exercise caution at this stage. Work quickly to prevent agar from solidifying. Also, position the organoids away from the edge of the column and keep them spaced out for optimal imaging results (**Figure 1B, magnified panel: organoids in agar column).** This can be achieved by pouring agar with a 200 µl pipette tip, gently moving the agar around the organoids with a needle, or simply rotating the syringe to shift the organoids within the column. Multiple organoids can be stacked in a single agar column.
27. Once the organoids are packed, place the syringe on a cooling pack to help the agar solidify. Gently rotate the syringe until the organoids are suspended in the agar, then let the agar harden.
28. After solidification, store the syringe at 4 ^°^C for at least 30 minutes to ensure the column is fully solidified. ▲ CRITICAL: Use a razor blade to make an angled cut at the distal end (away from the organoids) to mark their position for imaging. This helps identify the samples more quickly under the microscope. **Sequential dehydration and tissue clearing (Figure 1B)** ▲ CAUTION: Conduct these steps in the protocol under the fume hood. Ethanol is flammable. Wear protective gear as ethanol can cause eye irritation.
29. Prepare a series of ethanol dilutions (90%, 70%, 50%, and 30%) by diluting absolute ethanol with distilled water.
30. The agar columns with embedded organoids can now be gently extruded from the syringe into 15 mL pre-labelled Falcon tubes containing 30% ethanol by pressing the piston.
31. Incubate in 12-14 ml of 30% ethanol on a rocking platform for 2-3 hours at room temperature.
32. Incubate in 12-14ml of 50% ethanol on the roller for 2-3h at room temperature.
33. Incubate in 12-14 ml of 70% ethanol on the roller for 2-3 hours at room temperature. Alternatively, the roller can be placed at 4 ^°^C, and incubation in 70% ethanol can be done overnight.
34. Incubate in 12-14ml of 90% ethanol on the roller for 4h at room temperature.
35. Incubate in 12-14 ml of absolute ethanol on the roller for 4 hours at room temperature. The roller can then be set to 4 ^°^C, and incubation in 100% ethanol can continue overnight.
36. The next day, remove as much remaining ethanol as possible.
37. Add 10 mL of ethyl cinnamate to the tubes and place them in a tube rack covered with aluminum foil. ▴ CRITICAL: The agar column becomes translucent at this point. Place the tubes so that the column inside each tube does not come into contact with moisture. This is because moisture can irreversibly make the column opaque, rendering it unusable for imaging.
38. Incubate the samples overnight in ethyl cinnamate without disturbing the column. The column then becomes transparent and sinks in the tube. After this, the sample is ready for imaging. ▪ PAUSE POINT Samples can be stored at room temperature in the dark in ethyl cinnamate for several months before imaging, but it is recommended to image them within the same week after clearing to prevent loss of sample quality. **Box 1: Preparation of assembly for sample mounting (Figure 1C)** Sample mounting for light sheet microscopy of cleared samples is an essential step in the LUCID-org workflow. This technique involves converting an insulin syringe, a needle, and sticky tape into a cost-effective, durable, and reusable custom assembly. The finished assembly is shown in **Figure 1C**. Procedure
39. First, prepare a custom-made needle hook by carefully bending a needle to form a hook. ▴ CAUTION: The needle poses a cutting hazard. Handle it carefully. Use a razor blade to carefully and cleanly cut away the tapering part of an insulin syringe to make it cylindrical. ▴ CAUTION: Using a razor blade poses a cutting hazard. Handle the razor blade carefully. Also, ensure the cut is clean to prevent damage to the sample column during mounting and unmounting.
40. Attach the long end of the needle to the outside of the flat-ended insulin syringe neatly with sticky tape. The gap between the needle tip and the syringe should be 5-8 mm. This hook supports the sample in space and keeps it in the correct position when inserted in Ethyl cinnamate (ECi) during imaging. ▴ CAUTION: When mounting, ensure the hook is not too large and does not brush against any optics in the microscope.
41. Clean a glass petri dish with distilled water, then rinse with 70% ethanol and absolute ethanol. Dry it by wiping with lint-free paper. ▴ CRITICAL STEP: Use only the glass petri dish; otherwise, it will be corroded/dissolved by Ethyl cinnamate.
42. Take the Falcon tube containing the cleared sample and carefully decant the Eci, along with the agar column, into the glass Petri dish.
43. Use metal forceps to gently lift the column and carefully insert it into the empty syringe. *▴* CRITICAL STEP During mounting, ensure the needle is turned to the ‘out’ position away from the agar column to prevent damage to the agar.
44. Once the agar column is inserted into the syringe, just turn the needle to the ‘in’ position.
45. Gradually move the piston to push the column out of the syringe and onto the hook so that it supports the column.
46. Next, the sample can be mounted in the sample holder and inserted into the microscope.
47. Proceed to step 39 of the main procedure. **Phase 4: Sample mounting and imaging (Figure 1C)** **Sample mounting:**
48. Start the LS7 system, the system PC, and the processing/storage PC.
49. Refractive index matching: Ensure the detection and illumination objectives are correctly installed. Also, adjust the correction rings on both objectives to match the refractive index of Ethyl Cinnamate (1.558). *▴* CRITICAL STEP: Properly adjusting the correction rings on both objectives is essential for achieving a clear light sheet. This significantly enhances imaging quality.
50. Mount the appropriate sample chamber for imaging correctly. See Box 1 for details.
51. Insert fresh Ethyl cinnamate using a 50-ml syringe and tubing into the chamber. It can be reused 3-4 times. Now the sample chamber is ready. ▴ CAUTION: Avoid spillage; immediately clean with absolute ethanol to prevent the corrosive action of Ethyl cinnamate.
52. Mount the syringe into the sample holder and place it into the microscope. **Light Sheet microscopy imaging** Use the system software to position the syringe so that the column is in front of the objective. ▴ CRITICAL: Here, we use the Carl Zeiss LS7 system and Zen Black software to operate the microscope. The steps described here will vary greatly depending on the user’s preferred microscope.
53. Open the 3-track setup for imaging. This setup has separate channels in different tracks, enabling individual adjustment of the lasers.
54. Locate the organoids using a combined brightfield and fluorescence overview at 0.36x magnification.
55. Once the organoids are located, you can adjust the laser power, exposure, magnification, field of view, detector size, and other settings. *Note*: Enable the Pivot Scan option during imaging. This mostly removes shadows that might be cast on the sample by artifacts such as dust particles stuck to the column.
56. Define a Z-stack and multi-view imaging (if needed), then capture images. Set the appropriate field size to encompass the entire organoid.
57. Repeat this for all the organoids in the column.
58. The images can be saved on the Swap drive on the processing PC.
59. Ensure the data is saved. Reset the syringe to the ‘Load position’, which moves the sample away from the objective for retrieval.
60. Gently remove the syringe from the holder and pull the agar column back into the syringe.
61. Move the needle hook away from the column and transfer the column into a properly labeled 15ml Falcon tube. Quickly refill the tube with ethyl cinnamate from the petri dish for storage. ▪ PAUSE POINT: Samples can be stored in Ethyl cinnamate in the dark at room temperature for months and imaged multiple times.
62. Close the software and turn off the system.
63. Retrieve the Ethyl cinnamate from the sample chamber using the attached syringe and tubing, then transfer it to a bottle for reuse.
64. Remove the sample chamber (when empty), the tubing, and the syringe, then rinse them with absolute ethanol.
65. Air-dry the chamber, syringe and the tubing before appropriate storage.

### Timing

Phases 1 and 2: Whole-mount immunostaining: ∼2 d Phase 3: Organoid embedding and tissue clearing: ∼3 d

Phase 4: Sample mounting and Light sheet microscopy: ∼2 d

## Results

### Generation of iPSC-derived human brain organoids and cytoarchitectural analysis using thin-sectioning and staining

We generated brain organoids from iPSCs using a modified version of our previously described methods^10,31^. This method is based on non-directed differentiation and excludes embryoid body formation as an intermediate step (**Figure 2A**). It enables consistent production of brain organoids with similar cytoarchitecture, overall size, morphology (**Figure 2A, bottom panel)**, and cellular diversity across multiple iPSC donors^31^. To analyze the organoid tissues, we sectioned Day 30 organoids at 10 µm thickness and performed immunostaining with specific antibodies. This enabled visualization of neural progenitor cell (NPC) organization, apicobasal polarity, and neuronal arrangement. Consistently, the organoids showed clear apicobasal polarity, with ARL13B-positive primary cilia located at the apical lumen of the ventricular zone (VZ), SOX2-positive NPCs arranged radially, and MAP2-positive differentiating neurons positioned basally (**Figure 2B, top**), TUJ1-positive early differentiating neurons forming a primitive cortical plate–like structure at this stage (**Figure 2B, middle and Supplementary Figure 1A**), and DCX-positive immature neurons at the VZs (**Figure 2B, bottom and Supplementary Figure 1B**). Quantification of key cytoarchitectural features, such as the number of ventricular zones, VZ thickness, and MAP2-positive neuronal layer thickness, across at least three independent batches demonstrated reproducibility of the differentiation protocol (**Figure 2C).** We used these brain organoids in the LUCID-org platform.

### The LUCID-org pipeline provides unbiased, quantifiable cytoarchitectural insights into brain organoids

Although thin-section imaging provides useful insights, it does not capture the full three-dimensional spatial organization of tissue. Therefore, we used the LUCID-org pipeline with whole-organoid immunostaining, embedding and tissue clearing with Ethyl Cinnamate, and custom mounting to visualize brain organoid cytoarchitecture in its intact 3D form using light-sheet microscopy (**Figure 1).** We observed high efficiency in immunostaining, with deep antibody penetration throughout the organoids (**Supplementary Figures 2A and 2B**). In our experiment, we describe the immunostaining of brain organoids using SOX2 (marks neural progenitors that define the ventricular zone), MAP2 (labels differentiating neurons on the basal side), and ARL13B (labels primary cilia extending into the ventricular lumen from progenitors) (**Supplementary videos 1 and 2**), although many other antibodies and probes are compatible with the pipeline (**Supplementary Figure 2B**). We outline the method step by step in the simplest way possible, emphasizing key steps tailored to the protocol. We used a Carl Zeiss Light Sheet 7 (LS7) microscope to image our processed organoids.

### Image acquisition of unsectioned whole-brain organoids

We imaged whole-brain organoids using a 5x low-magnification objective, which allowed us to capture the entire organoid volume within 3–5 minutes while covering its full depth. We performed image acquisition with Zen Black software. First, we positioned the agar column containing embedded organoids in front of the objective lens. We then identified the organoids using any fluorescence channel and placed them within the acquisition area. Next, we adjusted individual laser parameters, including laser intensity, exposure time, and offset. Finally, we acquired raw image stacks of either whole organoids or specific regions of interest.

### Image pre-processing

Channel misalignment is an inherent result of the optics in the light-sheet microscope and must be corrected to prevent artefacts. Raw image stacks were individually adjusted for channel alignment using Zen Black. The ‘drift’ in each channel was manually calculated (for example, around 30 steps) and then corrected, with one channel designated as the reference (for example, inputting 30 steps for the misaligned channel).

### Image acquisition: Whole organoid

Low-magnification imaging with the 5x objective, combined with a volumetric projection of the full organoid stacks, already provided valuable insights into overall morphology, including apicobasal organization, ventricular-zone–like progenitor domains, and the emergence of a primitive cortical plate on the basal side (**Supplementary videos 3 and 4**, **Supplementary Figure 2A**).

### Image acquisition: Selected regions

To better illustrate the cytoarchitecture of the brain organoids, we further focused on the neurogenic niches, the ventricular zones (VZ). To fully utilize the capabilities of light-sheet microscopy and gain deeper insights into the organization of brain organoids, we performed volumetric imaging with a 20X objective and 2X zoom. Using this setup, we captured a selected VZ spanning approximately 200 µm in tissue depth within 2-4 minutes of acquisition (**Figure 3A, top panel,** and **Supplementary video 5**).

**Figure 3:**
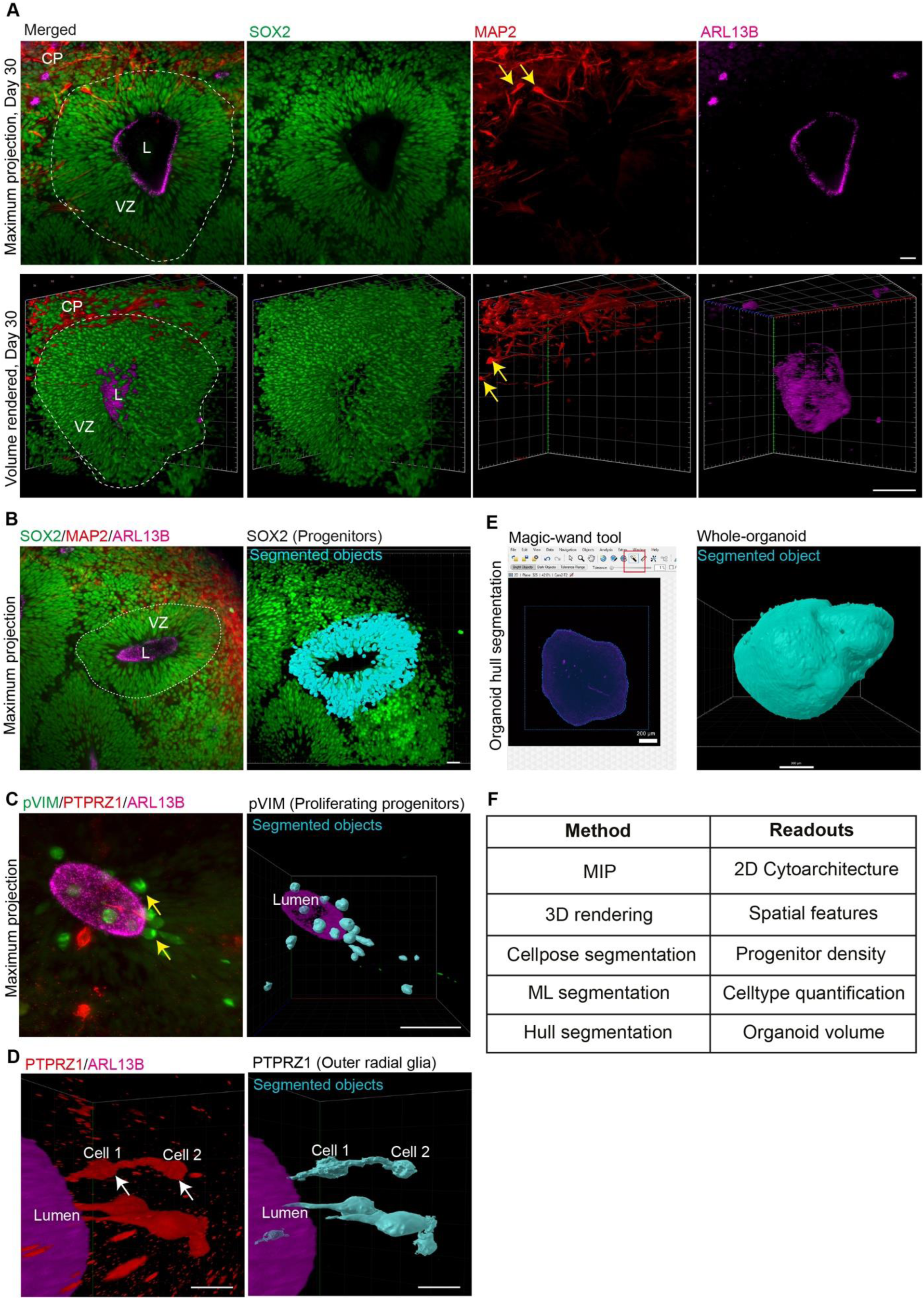
Image acquisition, visualization, and segmentation tools for whole mount imaging and quantification of cytoarchitectural features of brain organoids **A.** Top panel, Maximum Intensity Projection (MIP) light sheet image of a representative VZ captured using a 20X objective with a zoom factor 2. Organoid stained with anti-SOX2 (NPCs, green), anti-MAP2 (Neurons, red), and anti-ARL13B (Cilia, magenta) antibodies. Yellow arrowheads indicate the cell bodies of MAP2-positive neurons indicative of immature neurons forming a primitive CP. Scale bar top panel, 20 μm. Bottom panel-3D volumetric rendering and visualization of the VZ from the top panel, highlighting spatial features like the VZ depth, neuronal distribution, and lumen morphology in a 3D space. Yellow arrowheads indicate the neuronal cell bodies. Scale bar, 50 μm. The associated video for this panel is Supplementary video 5. **B.** Output of the Progenitor Segmentation Pipeline based on the Cellpose algorithm applied to the SOX2-positive progenitors in the VZ. Segmentation masks for the progenitors (right panel) overlap with the staining signal. Scale bar, 20 μm. The associated videos for this panel are Supplementary videos 16 and 17. **C.** Output of the Machine Learning Segmentation pipeline based on trained models to segment actively proliferating progenitors (pVIM-positive cells, green) surrounding the VZ at the lumen (ARL13B, magenta). The segmentation masks (right, cyan) overlap with the staining signal indicated by arrowheads (MIP image, left). Scale bar, 50 μm. **D.** Output of the Machine Learning Segmentation pipeline based on trained models to segment outer radial glia (PTPRZ1-positive cells, red) that radially migrate from apical to basal at the vicinity of the VZ lumen (ARL13B, magenta). The segmentation masks (right, cyan) recapitulate the morphologies of the radial glia indicated by white arrowheads (MIP image, left). Scale bar, 10 μm. The associated videos for this panel are Supplementary videos 12 and 13. **E.** ‘Magic-wand’ based automatic detection of macroscopic features, such as organoid surface, on whole-organoid stacks captured with the 5x objective. It was used to segment entire organoids for quantifying organoid hull volumes. Scale bar, 200 μm. **F.** Table summarizing the methods applied and their respective readouts.

Although light-sheet microscopy may not achieve the same spatial resolution as traditional confocal imaging, it provides unprecedented quantitative and volumetric insights into organoid cytoarchitecture. Imaging Day 30 brain organoids revealed a highly organized group of SOX2-positive (green) neural progenitors (NPCs) with characteristic palisade-like shapes surrounding the ventricular lumen (**Figure 3A**). The lumen itself was outlined by ARL13B-positive (magenta) primary cilia extending into the ventricular space. Basal to the ventricular lumen, MAP2-positive differentiating neurons were visible (**Supplementary video 5**). Importantly, in addition to neuronal processes, MAP2 staining was localized to neuronal cell bodies (yellow arrows), indicating that the organoids remain in an immature developmental stage at this point (**Figure 3A**).

### Quantifications

Light-sheet microscopy enabled unbiased quantification of features from the acquired volumetric datasets. Furthermore, the use of segmentation algorithms such as Cellpose, Blob finder, and Machine Learning could extract these features with high confidence and minimal manual intervention. By introducing the volumetric datasets into our custom data processing pipelines in Arivis Pro, we quantified structural parameters, including the number of SOX2-positive NPCs in a given volume (progenitor density), the number of phospho-Vimentin-positive apically positioned progenitors, PTPRZ1-positive outer radial glial cells migrating from apical to basal, the total organoid volumes, lumen numbers in organoids, and lumen volumes.

### 3D rendering

3D rendering is essential for visualizing and understanding the spatial organization of brain organoids, capturing a level of detail often missing in traditional imaging methods. To convert two-dimensional data into three-dimensional models, we rendered the entire dataset volumetrically. We utilized either the full stack depth across all VZs or selected data subsets after preprocessing. Volume reconstruction was carried out using Arivis Pro or Zen Blue software, as specified. This approach allowed us to visualize the ventricular zone as a continuous 3D structure and to examine the spatial distribution of neural progenitors and differentiating neurons within the organoid. Notably, primary cilia around the ventricular lumen formed a pocket-like structure (**Figure 3A, bottom panel**).

### Segmentation: Progenitor segmentation pipeline

To quantify progenitor cell abundance in the VZs, we used our custom progenitor segmentation pipeline **(Supplementary Figures 3A and 3B**). We first trimmed the dataset with a tailored view, then denoised it to enhance the signal-to-noise ratio. Next, we segmented neural progenitor cells (NPCs) using the deep-learning algorithm Cellpose, utilizing SOX2 staining as input. We iteratively refined the segmentation parameters with an interactive preview to quickly validate and optimize the results. The pipeline produced segmentation masks (turquoise) representing progenitor cells that closely overlapped with the SOX2 staining dataset. We then extracted these segmented features as objects for further data cleaning and quantitative analysis. Additionally, we developed a metric called ‘progenitor density’ to objectively measure and characterize the progenitor population within the VZs. This metric accounted for variations in VZ size within and between organoids.

To measure progenitor density, we created a volumetric data subset covering 25 steps upstream and downstream of the average lumen position, using a 20X objective lens with a zoom factor of 2 and a step size of 0.350 µm. This generated a z-stack of 50 steps, representing an imaging depth of approximately 17.5 µm. We consistently used this imaging setup across all acquisitions and subsets.

To limit quantification to the ventricular zone (VZ) and exclude extraventricular progenitors, we masked objects outside the VZ prior to segmentation. Then, we applied the Progenitor Segmentation pipeline to these datasets to determine progenitor density, defined as the number of progenitor cells within the analyzed volume. Progenitor density accurately reflects the biological structure of the VZs in brain organoids. This analysis revealed the spatial distribution of SOX2-positive neural progenitors within the VZ and enabled unbiased quantification of progenitor cells (**Figure 3B).**

### Segmentation: Machine learning segmentation pipeline

Certain features, such as cell morphologies and their spatial distribution, are crucial for understanding disease mechanisms, but standard imaging methods often miss this spatial information. To fix this, we created a custom machine-learning segmentation pipeline **(Supplementary Figure 4A-C)** to quantitatively extract cell-type–specific features from volumetric light-sheet datasets. We first pre-processed the channel-aligned data stacks. Then, we trained the machine-learning segmentation tool to identify specific features, such as Phospho-Vimentin (pVIM)-positive proliferating apical progenitor cells ^3^ ^32^, by manually annotating signal and background segments in a few sections. After training, we applied the model to the entire pre-processed 3D volumetric dataset, generating segmentation masks that closely matched the staining data. We then refined the segmented objects based on size and shape to remove artifacts and exported the cleaned objects for further analysis.

We expanded this method to include additional cell populations, such as phospho-Vimentin (pVIM)–positive proliferating apical progenitors, which line the ventricular surface marked by ARL13B-positive primary cilia (**Figure 3C**), and PTPRZ1-positive outer radial glia, which migrate radially from the apical to basal compartments ^33,34^ (**Figure 3D**). This workflow revealed the three-dimensional spatial arrangement of these distinct progenitor groups and enabled unbiased, volume-wide measurement of their distribution. Lastly, we used a ‘magic-wand’ tool (for lasso-like selection) to segment the organoid hull for whole organoid volume measurements (**Figure 3E**). Methods and their respective readouts are summarized in **Figure 3F**.

### LUCID-org facilitates modeling of microcephaly caused by a *CENPJ* mutation

Having optimized for imaging and quantification, we plan to validate our developed methodologies in disease modeling. To do this, we generated brain organoids exhibiting microcephaly phenotypes and applied our LUCID-org pipeline. *CENPJ* (CPAP), a centrosomal gene, is mutated in patients with Seckel syndrome, a developmental disorder also characterized by severe microcephaly ^10^. We previously revealed disease mechanisms using brain organoids derived from Seckel patient cells and observed depletion of symmetrically proliferating neural progenitors. Here, we tested whether our optimized light-sheet imaging pipeline could validate and sensitively detect developmental defects associated with microcephaly.

Compared with healthy controls (brain organoids derived from IMR90 iPSCs), brain organoids from a *CENPJ*-mutant patient were significantly smaller, indicating an early developmental defect (**Figure 4A**). We also assessed organoid volumes using our machine-learning segmentation pipeline, which showed that *CENPJ*-mutant brain organoids had considerably smaller volumes than control organoids (**Figure 4B**). To explore the developmental plasticity of brain organoids, we analyzed Day 15 and 30-old organoids from healthy and microcephaly groups. At Day 15, control organoids displayed well-organized apicobasal polarity, with SOX2-positive neural progenitor cells (NPCs) forming a neural rosette around a ventricular lumen, and ARL13B-positive primary cilia localized to the apical surface (**Figure 4C, top panel)**. Conversely, microcephaly organoids showed a significant disruption of this organization: NPCs were scattered randomly and rarely formed a clear neural rosette (**Fig 4C, bottom panel**). Control organoids contained only a few MAP2-positive differentiating neurons, indicating they remained in an actively proliferative state. Microcephaly organoids at this stage already showed neurons randomly dispersed along the apical–basal axis, suggesting that progenitors are unstable in maintaining their proliferative capacity (**Figure 4C**). By Day 30, control organoids exhibited a highly organized, palisade-like structure of SOX2-positive progenitors, which was severely distorted in *CENPJ*-mutant organoids (**Figure 4D, Supplementary videos 6-9)**. Likewise, the MAP2-positive region, representing the developing cortical plate, formed a distinct neuronal layer in control organoids, while MAP2-positive neurons were randomly spread through the tissue in *CENPJ*-microcephaly organoids (**Figure 4D, Supplementary videos 10 and 11)**. Our imaging revealed numerous smaller micro-rosettes and a disrupted neuronal pattern in microcephaly organoids (**Figure 4D**). Importantly, these abnormalities were visible in intact three-dimensional volumetric datasets, providing spatial information that is largely inaccessible with traditional thin-sectioning or confocal microscopy (**Figure 4E, Supplementary Figure 5)**. Our analysis also showed reduced VZ thickness (**Figure 4F**) and disrupted MAP2-positive region (**Figure 4G**) in *CENPJ*-microcephaly organoids. Overall, these findings (summarized in **Figure 4H**) demonstrate that our fixation, clearing, and staining workflow preserves the native cytoarchitecture of brain organoids and enables reliable detection of structural defects relevant to microcephaly caused by the *CENPJ* mutation.

**Figure 4:**
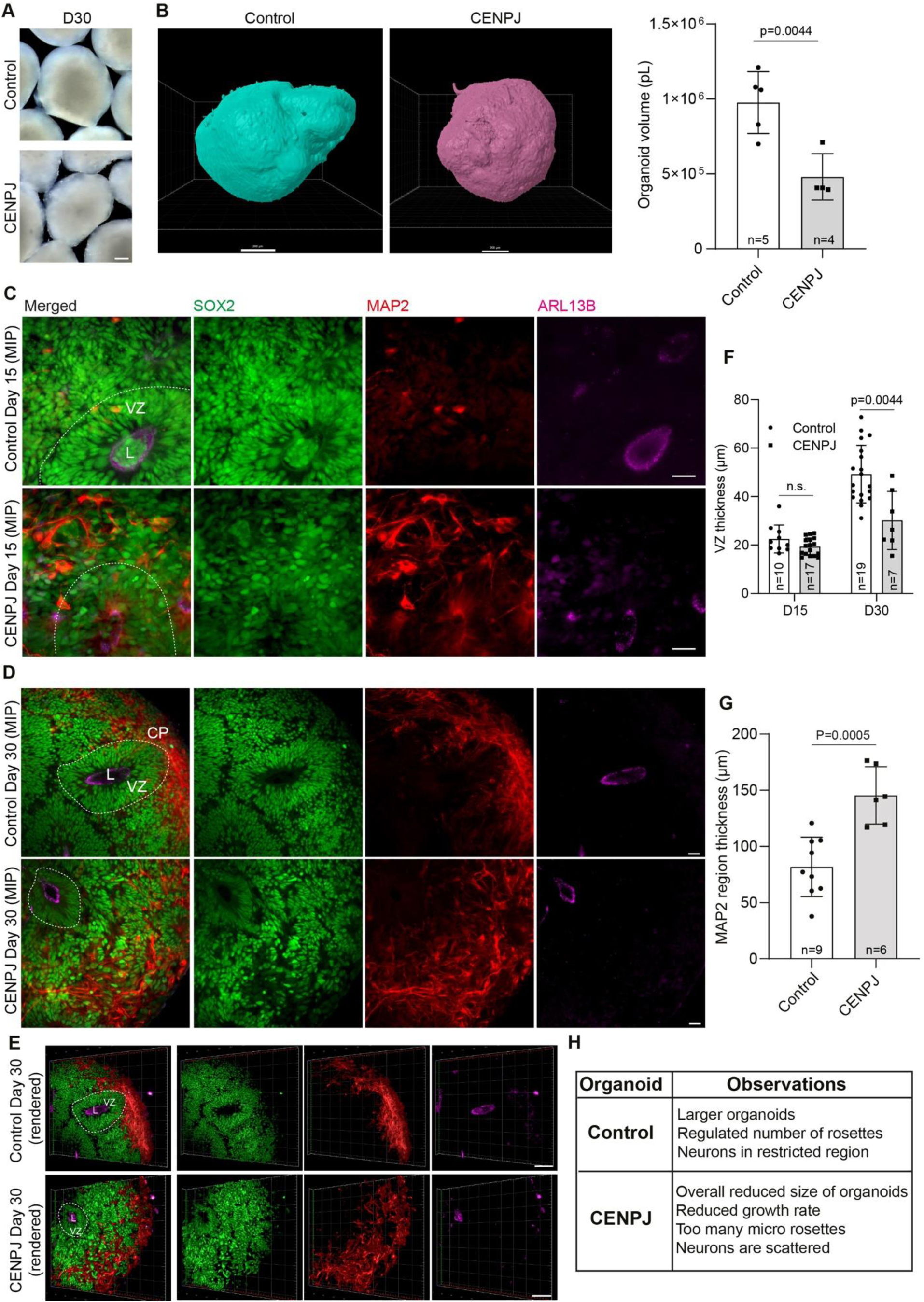
LS imaging of control and *CENPJ*-microcephaly organoids quantifies developmental defects. **A.** Macroscopic images (left) of control and *CENPJ*-microcephaly organoids at D30. Scale bar, 200 µm. **B.** Representative segmented whole-organoids from control and *CENPJ*-microcephaly organoids. The bar diagram at the right quantifies their volumes. The analysis was performed on n=5 (control) and n=4 (CENPJ) organoids across two independent batches. The values for “n” and “p” are indicated in the plot. Error bars represent standard deviation. Student’s t-test was used for statistical analysis. Scale bar, 200 µm. **C.** MIP image of representative VZ of control organoids (top panel) and *CENPJ*-microcephaly organoids (bottom panel) at D15. SOX2 specifies NPCs (green). MAP2 marks differentiating neurons (red). ARL13B (magenta) marks primary cilia lining the VZ lumen. Scale bar, 20 μm**. D.** MIP image of representative VZ of control organoids (top panel) and *CENPJ*-microcephaly organoids (bottom panel) at D30. SOX2 specifies NPCs (green). MAP2 marks differentiating neurons (red). ARL13B (magenta) marks primary cilia lining the VZ lumen. Scale bar, 20 μm. The associated videos for this panel are Supplementary videos 6-11. **E.** 3D volume rendered sections of representative images of VZs from control (top panel) and *CENPJ*-microcephaly (bottom panel) brain organoids at D30, highlighting their spatial organization. SOX2 specifies NPCs (green). MAP2 marks differentiating neurons (red). ARL13B (magenta) marks primary cilia lining the VZ lumen. Scale bar, 50 μm. **F.** Quantification of VZ thickness for control and *CENPJ*-microcephaly brain organoids at D15 and D30. Error bars represent standard deviation. The analysis was performed on n=10 (control) and n=17 (*CENPJ*) VZs for Day 15, and on n=19 (control) and n=7 (*CENPJ*) VZs across two independent batches of organoids. The values for “n” and “p” are indicated in the plot. Student’s t-test was used for each time point for statistical analysis. **G.** Quantification of MAP2 region thickness (neuronal distribution) in control and *CENPJ*-microcephaly brain organoids at D30. Error bars represent standard deviation. The analysis was performed on n=9 (control) and n=6 (*CENPJ*) regions across two independent batches of organoids. The values for “n” and “p” are indicated in the plot. Student’s t-test was used for statistical analysis. **H.** Table summarizing key observations from volumetric imaging of control and *CENPJ*-microcephaly brain organoids.

Next, we employed our Machine Learning Segmentation pipeline to quantitatively extract cell-type–specific features impacted in microcephaly (**Supplementary Figure 4A-C**). Specifically, we examined Phospho-Vimentin–positive apical progenitors and PTPRZ1-positive outer radial glial (oRG) cells. Apical progenitors typically divide symmetrically at the ventricular surface to expand the progenitor pool, while oRG cells are a specialized class of NPCs that migrate from the apical to basal axis to form the outer subventricular zone (oSVZ), a process essential for neocortical development. Using our segmentation pipeline, we consistently identified oRG cells based on their distinctive elongated morphology.

In healthy control brain organoids, oRG cells were abundant and exhibited the expected polarized morphology. In contrast, microcephaly organoids showed a significant reduction in oRG cells, indicating these cells are produced in insufficient numbers from apical progenitors (**Figure 5A, Supplementary videos 12 and 13)**. We then focused on Phospho-Vimentin–positive progenitors in the apical region. Phospho-Vimentin typically marks dividing progenitors in this area. Our pipeline’s three-dimensional segmentation allowed visualization of these cells within the intact organoid structure and enabled robust quantification in both conditions. Interestingly, microcephaly organoids contained more Phospho-Vimentin–positive cells than controls (**Figure 5B and Supplementary Figure 6)**, but these cells were randomly scattered, suggesting they are stalled or arrested at the ventricular surface and may undergo premature differentiation directly into neurons. This interpretation aligns with the widespread presence of neurons throughout the organoid volume and the polarity defects observed in microcephaly samples.

**Figure 5:**
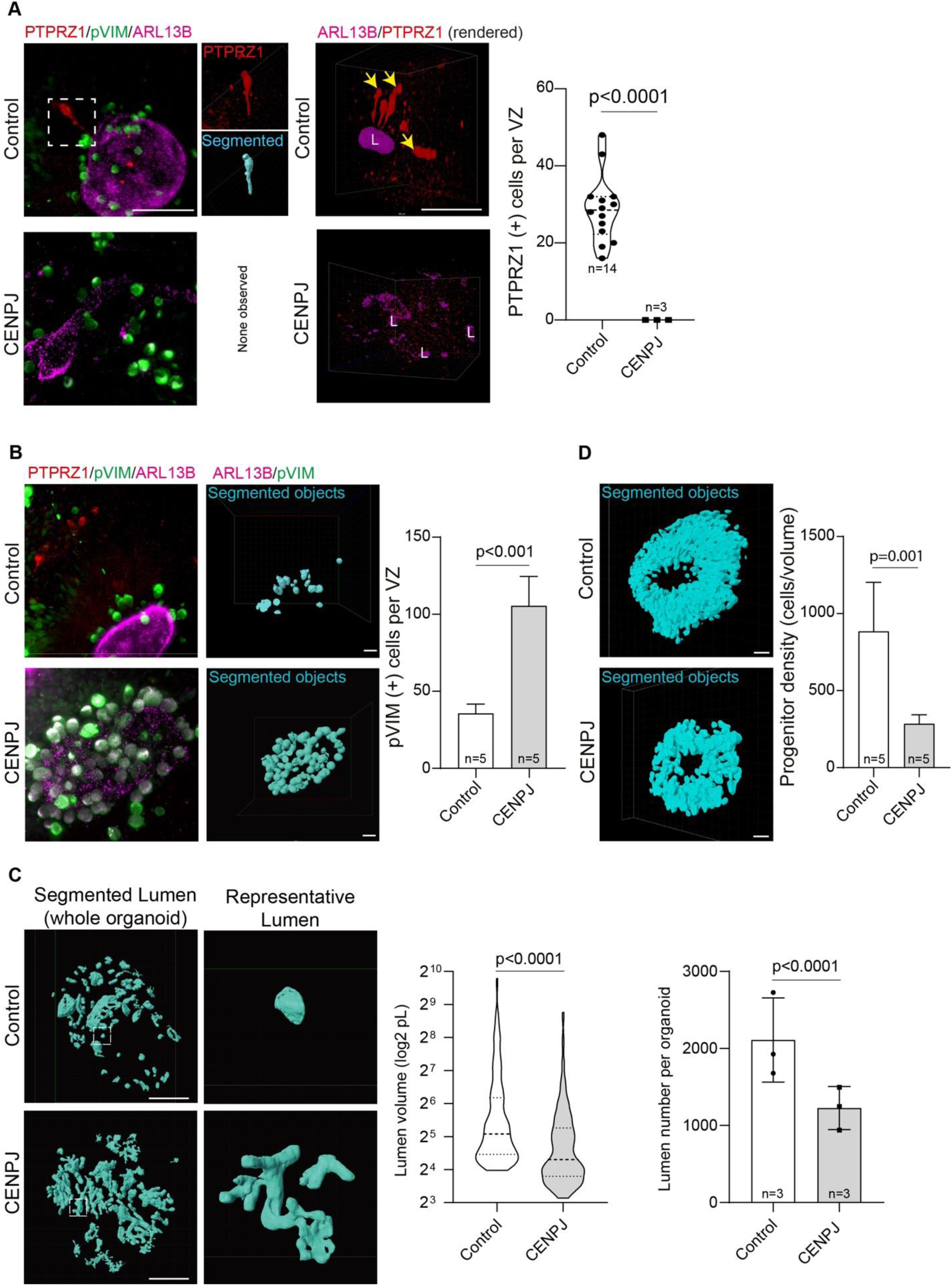
Quantification of 3D spatial features from volumetric LS imaging data highlights altered cellular dynamics and architectural defects in *CENPJ*-microcephaly brain organoids. **A.** MIP image of representative VZs from control (top panel) and *CENPJ*-microcephaly organoids (bottom panel) stained with anti-PTPRZ1 (outer radial glia, red), anti-phospho-Vimentin (proliferating progenitors, green), and anti-ARL13B (Cilia, magenta). Inset highlights an outer radial glial cell and the output from the Machine Learning Segmentation pipeline (cyan) trained to segment outer radial glial cells based on PTPRZ1 staining signal. The segmentation mask overlays tightly onto the staining signal. The panel on the right shows 3D volumetric renderings of the respective Lumen. Yellow arrowheads indicate the outer radial glia near the lumen (magenta). Scale bar, 200 µm. The graph on the right quantifies the number of PTPRZ1-positive cells per VZ in control and *CENPJ*-microcephaly brain organoids. The analysis was performed on n=14 (control) and n=3 (*CENPJ*) whole-VZ stacks from two independent organoid batches. The values for “n” and “p” are indicated in the plot. Student’s t-test was used for statistical analysis. **B.** MIP image of representative VZs with the same staining as in **A,** with output from the Machine Learning Segmentation pipeline trained to segment proliferating progenitors based on phospho-Vimentin staining signal. The segmentation masks overlay tightly onto the staining signal. Scale bar, 20 µm. The graph on the right quantifies the number of pVIM-positive cells per VZ in control and *CENPJ*-microcephaly organoids. The analysis was performed on n=5 (control) and n=5 (*CENPJ*) whole-VZ stacks across two independent batches of organoids. The values for “n” and “p” are indicated in the plot. Student’s t-test was used for statistical analysis. **C.** Output of Machine Learning Segmentation pipeline trained for segmentation of lumen based on ARL13B staining signal in whole organoids for control (top panel) and *CENPJ*-microcephaly (bottom panel). Insets highlight the morphologies of representative isolated lumen from control and *CENPJ*-microcephaly organoids. Scale bar, 200 μm. The graphs show quantification of lumen volume from the 100 largest lumens in each sample (left) and total number of lumens (right) for control and *CENPJ*-microcephaly organoids. Error bars represent standard deviation. The analysis was performed on n=3 (control) and n=3 (*CENPJ*) organoids across two independent batches of organoids. The values for “n” and “p” are indicated in the plot. Student’s t-test was used for statistical analysis. **D.** Output of the machine learning segmentation pipeline trained to identify progenitors at a specific depth in the VZ of control and *CENPJ*-microcephaly organoids. The segmented objects represent SOX2-positive progenitors. Scale bar, 20 μm. The graph on the right shows progenitor density (number of progenitors at a given VZ depth) per VZ in control and *CENPJ*-microcephaly organoids. Error bars indicate standard deviation. The analysis involved n=5 (control) and n=5 (*CENPJ*) whole-VZ subsets from two independent organoid batches. The values for “n” and “p” are indicated in the plot. Student’s t-test was used for statistical analysis.

Similarly, our segmentation analysis enabled both qualitative and quantitative assessments of ventricular lumen architecture. Lumen was more branched, highly convoluted, and appeared connected like a network in microcephaly organoids, whereas they formed distinct and better-organized units in the control organoids (**Figure 5C, right panel and Supplementary videos 14 and 15**). Additionally, the estimated ventricular lumen volume and overall lumen numbers were significantly smaller in *CENPJ*-microcephaly organoids (**Figure 5C**). These findings were consistent with the presence of multiple micro-rosettes in microcephaly organoids (**Figure 5C**). Furthermore, we used our progenitor segmentation pipeline (**Supplementary Figure 3A-B**) on data subsets from Control and *CENPJ*-mutant brain organoids to evaluate progenitor densities. Notably, it showed that progenitor densities in the VZs of *CENPJ*-mutant brain organoids were significantly lower than in their control counterparts (**Figure 5D, Supplementary videos 16 and 17**). This suggests an insufficient number of progenitors, possibly due to premature differentiation in *CENPJ*-mutant brain organoids, a typical feature of microcephaly as previously described^10^.

## Discussion

In this study, we optimized light-sheet imaging and machine-learning–based three-dimensional segmentation to establish a protocol for quantitative imaging of brain organoids. These methods are designed to resolve the cytoarchitecture of brain organoids in their intact state, thereby avoiding the loss of spatial information inherent in invasive thin-sectioning approaches. Although the optimized workflow may require special expertise, it provides substantially improved structural information in a fully quantitative and unbiased manner, enabling more rigorous functional and phenotypic analyses (**Figures 1-5).**

To experimentally validate our approach, we used *CENPJ*-mutant brain organoids as a disease model and observed cytoarchitectural defects that recapitulate key neurodevelopmental abnormalities in patients with microcephaly (**Figure 4-5**).

In particular, our LUCID-org approach offered the following observations in analyzing the mutant organoids, which were not readily derived from conventional sectioning and imaging: ***i)*** Disruptions in progenitor organization in volumetric terms, ***ii)*** 3D VZ architecture, ***iii)*** fragmented ventricular structures, ***iv)*** presence of multiple micro-rosettes rather than a single, well-organized VZ lumen.

Volumetric analysis further revealed several phenotypes indicative of developmental disorders in *CENPJ* brain organoids. For instance, despite comparable total lumen volume, the increased number of smaller, more convoluted lumens in the mutant brain organoid indicates impaired ventricular zone integrity, consistent with defects in the symmetric expansion of apical progenitors. These architectural abnormalities correlated with altered progenitor dynamics, including an unexpected accumulation of Phospho-Vimentin–positive apical progenitors at the ventricular surface, suggesting stalled or aberrant mitotic progression. Concomitantly, we observed a marked reduction in PTPRZ1-positive outer radial glial cells and a loss of spatially restricted neuronal layers, with MAP2-positive neurons distributed throughout the organoid volume. Together, these findings highlight the strength of intact three-dimensional analysis to resolve lineage and spatial defects that underlie impaired tissue growth (**Figure 5**).

Despite several limitations, the described approach provides an experimentally adaptable and unbiased strategy to link genetic perturbations to multiscale structural phenotypes in human brain organoids. We anticipate that this methodology will be useful for other organoid systems and disease models, possibly enabling image-based, quantitative disease modeling of human neurodevelopmental disorders.

## Acknowledgement

We thank the members of the Laboratory of Centrosome and Cytoskeleton Biology (CCB). J.G. acknowledges support from Innovation funding from BMBF VIP+ (03VP10540), and Deutsche Forschungsgemeinschaft (DFG, German Research Foundation)—Project-ID 503306912—FOR5547; DFG; O.S.V. is supported by the Medical Scientist Program, IZKF, University Hospital Jena. The authors acknowledge Angnes von Keller and Maurizio Abbate from Zeiss for the introductory training and support for image acquisition and processing, respectively.

## Author contributions

O.S.V. developed the protocols for organoid generation, immunostaining, embedding, sample mounting, image acquisition, processing, and analysis pipelines for 3D quantifications in Arivis Pro and the LUCID-ORG workflow. A.V.J. performed brain organoid generation, immunostainings, image acquisition, and quantifications. H.A.O. provided support with protocol standardization. E.G. provided the *CENPJ*-microcephaly patient iPSC cells for organoid generation. J.G. supervised the project. O.S.V., A.V.J., and J.G. wrote the manuscript. All authors read and contributed to its revision.

## Supplementary videos

**Video 1.** Z-series subset of a representative VZ from control organoid stained with anti-SOX2 (progenitors, green), anti-MAP2 (neurons, red), and anti-ARL13B (cilia, magenta).

**Video 2.** 3D volumetric rendering for the Z-series subset of a representative VZ from control organoid stained with anti-SOX2 (progenitors, green), anti-MAP2 (neurons, red), and anti-ARL13B (cilia, magenta).

**Video 3.** 3D volumetric projection of a representative control whole organoid stained with anti-SOX2 (progenitors, green), anti-MAP2 (neurons, red), and anti-ARL13B (cilia, magenta).

**Video 4.** 3D volumetric projection of a representative *CENPJ*-microcephaly whole organoid stained with anti-SOX2 (progenitors, green), anti-MAP2 (neurons, red), and anti-ARL13B (cilia, magenta).

**Video 5.** 3D volumetric rendering of the full Z-series of a representative VZ from control organoid stained with anti-SOX2 (progenitors, green), anti-MAP2 (neurons, red), and anti-ARL13B (cilia, magenta).

**Video 6.** 3D volumetric rendering of a Z-series subset of a representative VZ from a control organoid stained with anti-SOX2 (progenitors, green), anti-MAP2 (neurons, red), and anti-ARL13B (cilia, magenta).

**Video** 7. Dataset from Video 6 highlighting progenitors (green) and lumen (magenta).

**Video 8.** 3D volumetric rendering of a Z-series subset of a representative VZ from *CENPJ*-microcephaly organoid stained with anti-SOX2 (progenitors, green), anti-MAP2 (neurons, red), and anti-ARL13B (cilia, magenta).

**Video 9.** Dataset from Video 8 highlighting progenitors (green) and lumen (magenta).

**Video 10.** Dataset from Video 6 highlighting neurons (red) and lumen (magenta).

**Video 11.** Dataset from **Video 8** highlighting neurons (red) and lumen (magenta).

**Video 12.** 3D volumetric rendering of the entire Z-series of a representative lumen from a control organoid, emphasizing PTPRZ1-positive outer radial glia (red) and lumen (magenta).

**Video 13.** 3D volumetric rendering of the entire Z-series of a representative lumen from *CENPJ*-microcephaly organoid, emphasizing PTPRZ1-positive outer radial glia (red) and lumen (magenta).

**Video 14.** Machine learning segmentation pipeline output highlighting the segmented lumen in a representative control whole organoid.

**Video 15.** Machine learning segmentation pipeline output highlighting the segmented lumen in a representative *CENPJ*-mutant whole organoid.

**Video 16.** Progenitor segmentation pipeline output highlighting progenitor density in a representative VZ from a control organoid.

**Video 17.** Progenitor segmentation pipeline output highlighting progenitor density in a representative VZ from a *CENPJ*-microcephaly organoid.

## Supplementary Figures

**Supplementary Figure 1.**
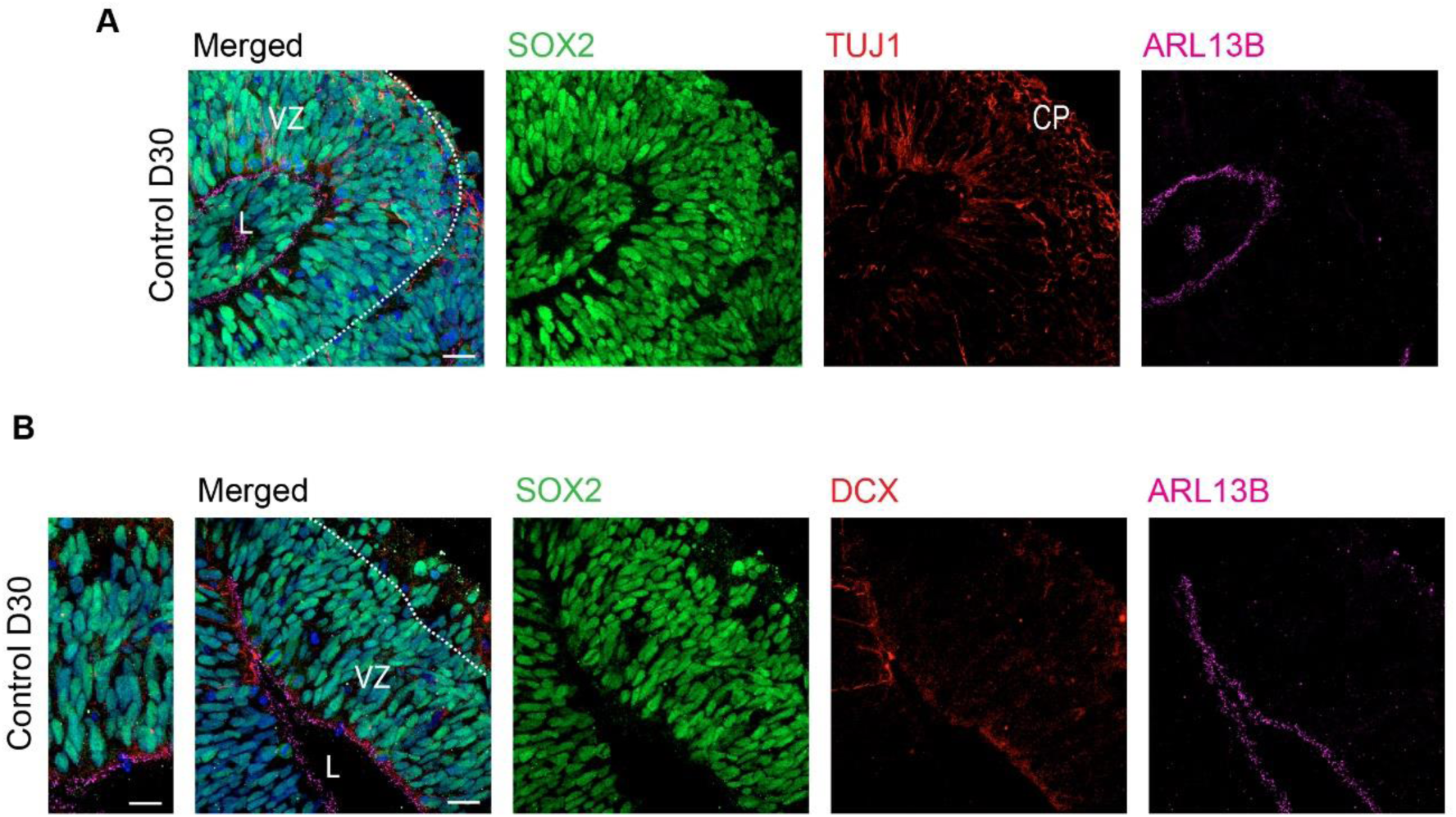
**A**. Immunofluorescence staining on cryosections of D30 control brain organoids stained with anti-SOX2 (progenitors, green), anti-TUJ1 (neurons, red), and anti-ARL13B (Cilia, magenta) antibodies, highlighting well-defined Ventricular Zones (VZs), Lumen (L), and the primitive Cortical Plate (CP) on the basal side. Scale bar, 20 μm. **B.** Similar staining as in **a,** with anti-DCX (immature neurons, red). Scale bar, 20 μm.

**Supplementary Figure 2.**
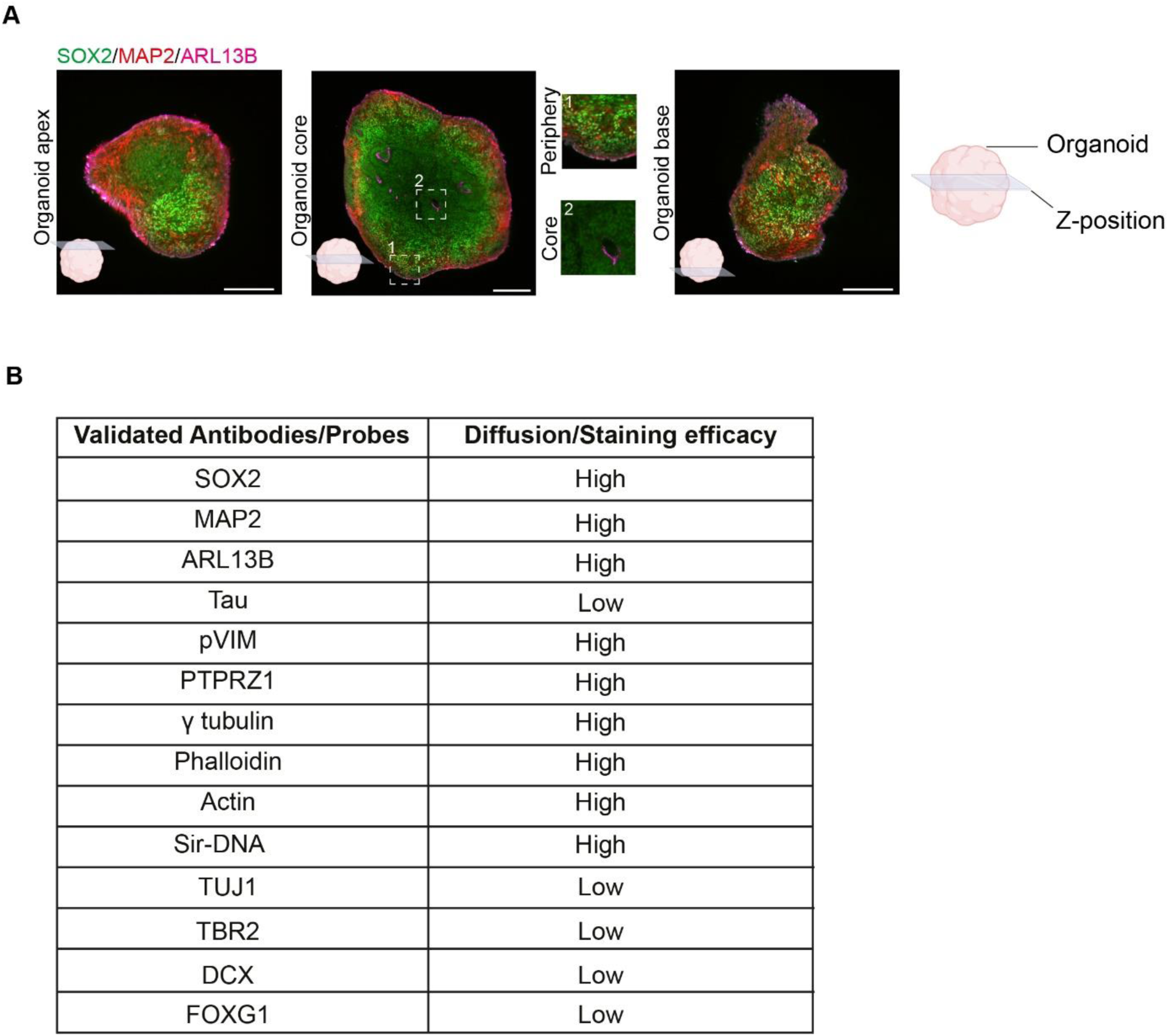
**A**. An optical section of a representative organoid stained with anti-SOX2 (progenitors, green), anti-MAP2 (neurons, red), and anti-ARL13B (cilia, magenta) at indicated Z-positions (illustrated as insets), highlighting the antibody penetration during whole-mount staining in LUCID-ORG workflow. The images represent the following: Left panel-organoid apex, middle panel-organoid core (inset 1-periphery and inset 2-core), and the right panel-organoid base with a legend for the illustration on the right. Scale bar, 200 μm. **B.** List of validated Antibodies and Probes and their diffusion/efficacies for staining in the LUCID-ORG workflow for light sheet microscopy.

**Supplementary Figure 3.**
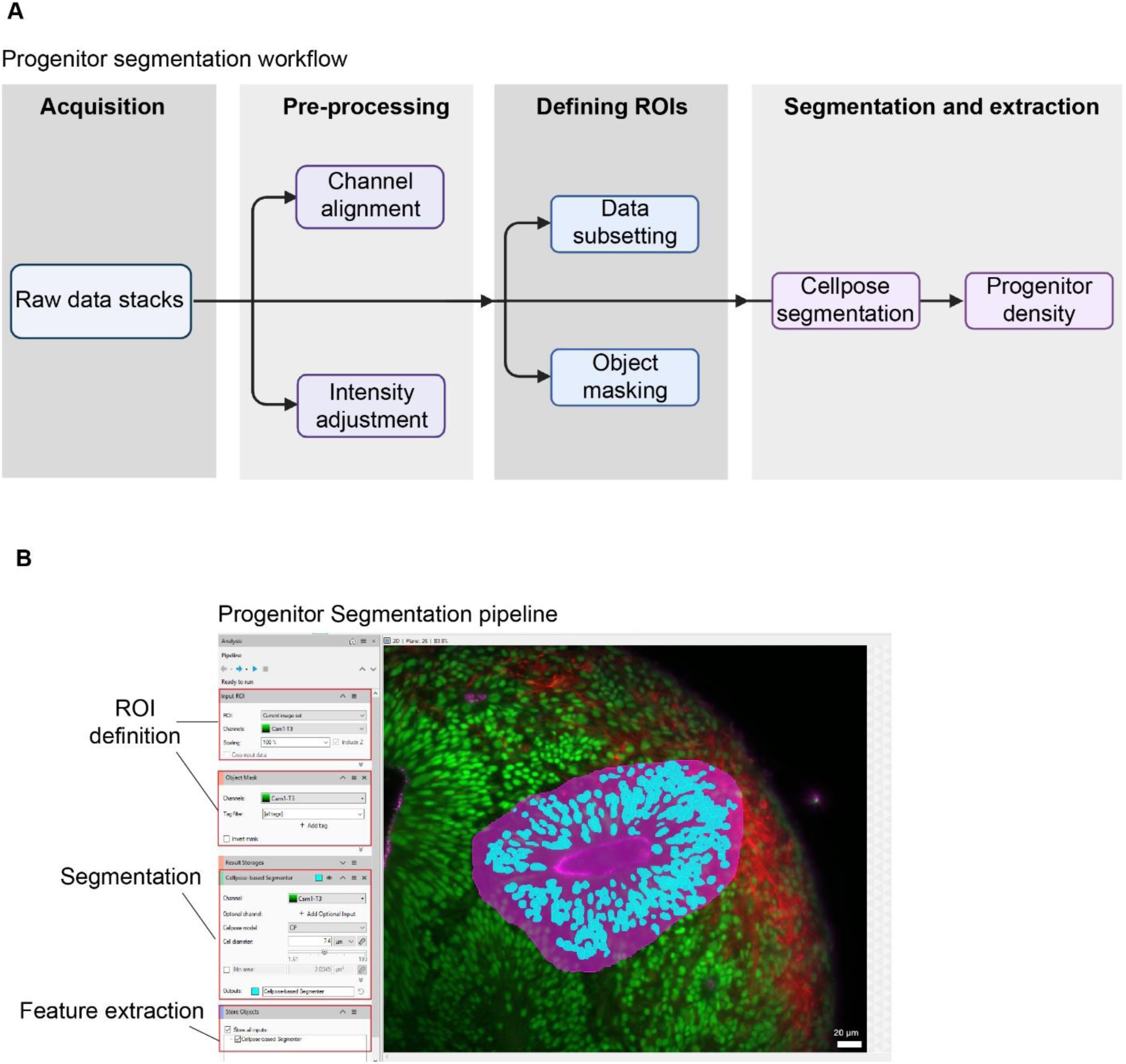
**A**. Overall workflow for Progenitor Segmentation using Cellpose algorithm. It comprises steps of data acquisition, preprocessing, ROI definition, and segmentation. **B**. Screenshot of the Progenitor Segmentation pipeline in Arivis Pro (insets 1 and 2-ROI definition, inset 3-Cellpose segmentation, inset 4-feature extraction). The image in the preview represents the pipeline’s output.

**Supplementary Figure 4.**
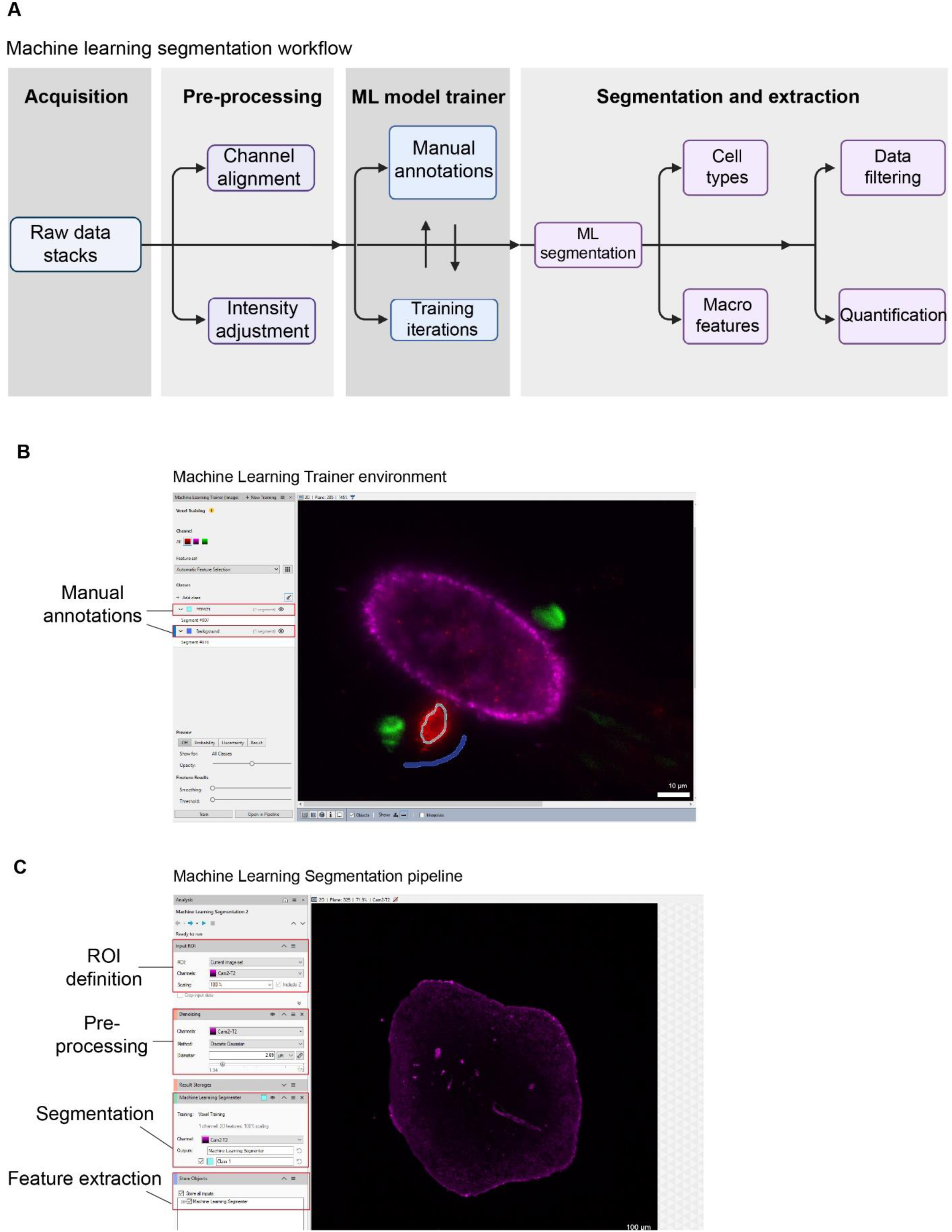
**A**. Overall workflow for Machine Learning Segmentation includes steps such as data acquisition, preprocessing, Machine Learning training, segmentation of cell types or macro features like lumen, followed by feature extraction. **B**. Screenshot of the Machine Learning Trainer environment in Arivis Pro. For model training, examples of manually annotated features, such as the PTPRZ1 staining signal (Cyan segment) and background (Blue segment), are shown. **C.** Screenshot of the Machine Learning segmentation pipeline in Arivis Pro (inset 1 - ROI definition, inset 2 - pre-processing, inset 3 - ML segmentation, inset 4 - feature extraction).

**Supplementary Figure 5.**
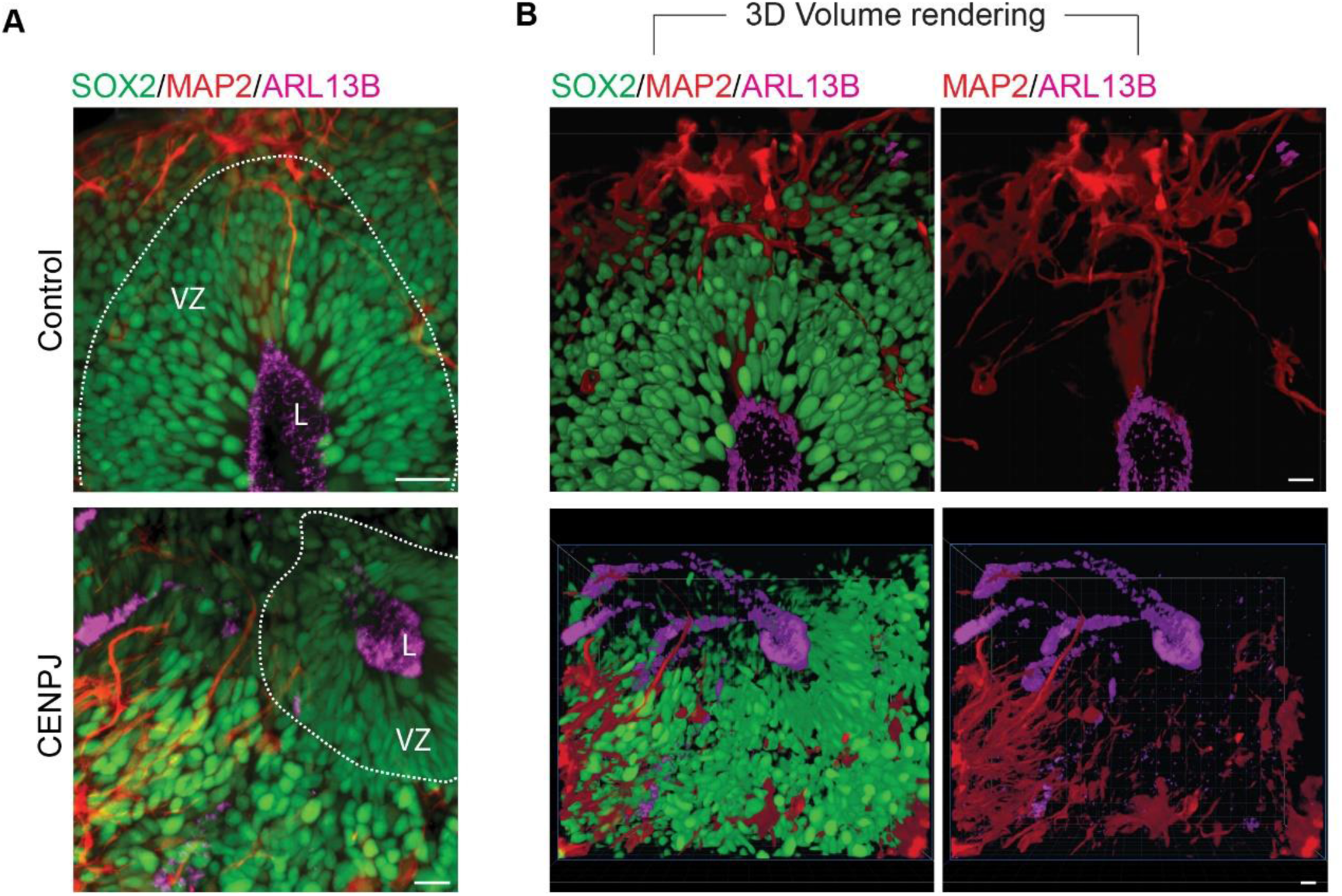
**A**. MIP image of representative VZ from control organoids (top panel) and *CENPJ*-microcephaly organoids (bottom panel), illustrating the spatial organization of progenitors, neurons, and lumen. The organoids were stained with anti-SOX2 (progenitors, green), anti-MAP2 (neurons, red), and anti-ARL13B (cilia, magenta). **B**. A 3D volumetric rendering of the data from **A**. Scale bar, 20 μm.

**Supplementary Figure 6.**
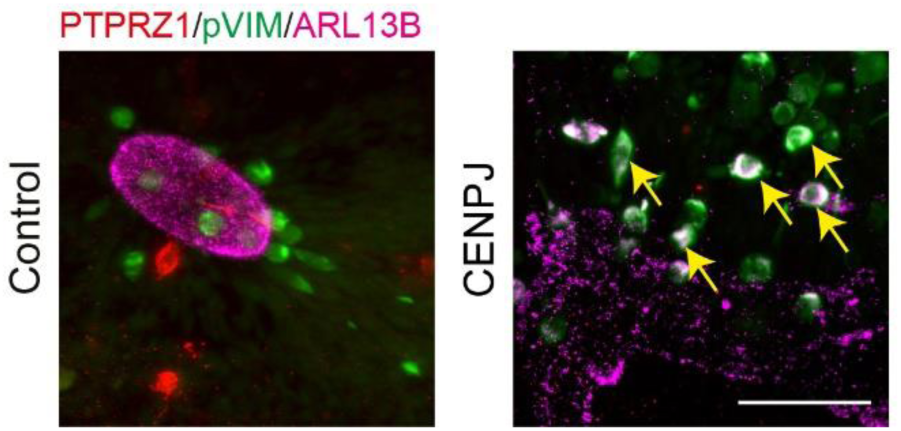
MIP image of a representative VZ from control and *CENPJ*-microcephaly organoids stained with anti-phospho-Vimenin (proliferating progenitors, green), anti-PTPRZ1 (outer radial glia, red) and anti-ARL13B (cilia, magenta). Yellow arrowheads highlight scattered pVIM-positive cells distant from the lumen surface in *CENPJ*-microcephaly organoids. Scale bar, 50 μm.

